# Pangenome and genome-scale essentiality in the emerging fungal pathogen *Candidozyma auris*

**DOI:** 10.64898/2026.05.12.724661

**Authors:** Joseph J Hale, Ajay J Larkin, Jackson R Rapala, Rebecca Hurto, Guolei Zhao, Brynn E Elson, Lydia Freddolino, Evan S. Snitkin, Teresa R O’Meara

**Affiliations:** Department of Microbiology and Immunology, University of Michigan Medical School; Ann Arbor MI 48019 USA; Gilbert S. Omenn Department of Computational Medicine and Bioinformatics, University of Michigan Medical School; Ann Arbor MI 48019 USA; Department of Biological Chemistry, University of Michigan Medical School; Ann Arbor MI 48019 USA; Program in Immunology, University of Michigan Medical School; Ann Arbor MI 48019 USA

**Author notes:** Correspondence to: Teresa O’Meara Evan Snitkin. denotes equal contribution (Joseph J Hale, Ajay J Larkin, Teresa R O’Meara).

## Abstract

*Candidozyma auris* is a multi-drug resistant fungal pathogen that causes hospital outbreaks, but genetic variation and gene function in this organism remain undercharacterized. Here, we performed pangenome analysis on 695 genetically representative isolates of *C. auris* and found that only 2.6% of gene families varied in their presence, mainly consisting of gene loss events or clade-specific genes, highlighting the clonal nature of these strains. Using microbial genome-wide association, we identified two loci associated with altered keratinocyte adherence, including genes with unknown function. To look broadly at gene function, we developed a genome-scale insertional mutagenesis approach and identified 614 high-confidence essential genes. Nearly one-third of these genes exhibited divergent essentiality compared to the model yeasts *Candida albicans* and *Saccharomyces cerevisiae*. Together, this study highlights organism-specific biology and provides a comprehensive resource for identifying species-specific determinants of virulence and targets for antifungal drug development.

## Introduction

*Candidozyma auris* is an often multi-drug resistant fungal pathogen that causes hospital-associated outbreaks ^1^. The mortality of systemic infections approaches 50%, and cases are on the rise due to the high colonization and transmission rates in healthcare settings. There are often few treatment options after systemic infection, and isolates resistant to at least two of the three commonly used classes of antifungals have been frequently observed ^2,3^. However, as a recently discovered organism, *C. auris* has a largely uncharacterized genome, an unknown set of essential genes, and poorly-understood genetic mechanisms for generating phenotypic variation. This lack of knowledge of essential pathways can limit our ability to identify potential new drug targets specific to this species. Additionally, our limited insight into the genetic changes experienced by lineages of *C. auris* can hinder our ability to predict the genotypic and phenotypic adaptations that occur in healthcare settings. Thus, there is a need to better understand the genetic variation that exists within and between the different clades of *C. auris*, as well as investigate the genes and pathways essential for its survival.

Pangenome analysis is an approach to identify the full collection of orthologous gene clusters across all the sequenced genomes for a given species, thus allowing an analysis of genetic diversity that cannot be captured in a single reference strain. Previous studies have shown that *C. auris* isolates tend to be highly clonal within each clade, with median pairwise SNP distances below 100 for clades I, III and IV ^4^. However, evidence for high rates of genomic rearrangement has also been found ^5–7^, implying that significant differences in gene content may exist between isolates and clades. Additionally, *C. auris* strains vary within and between clades in several clinically relevant phenotypes, such as antifungal resistance, adherence to skin and abiotic surfaces, and nutrient utilization ^1,8–11^. Hospital pathogen surveillance has produced a large number of sequenced isolates from diverse clades, time points, and geographic locations ^12–15^. Thus, a pangenome analysis of *C. auris* that takes its clonal nature and population structure into account has the potential to reveal the genetic changes experienced by hospital-associated isolates and lineages, in addition to identifying significant and clade-defining genetic variants for future studies.

Beyond presence and absence, the set of essential genes (those that are required for viability in non-stress conditions) are also uncharacterized in *C. auris*. One approach for defining the essential gene set is via insertional mutagenesis ^16^. In *C. auris,* the *piggyBac* transposon system was used to identify lncRNA that controls filamentation and *CDT1*, a calcium-calcineurin signaling pathway-sensitive ATPase that controls fluconazole tolerance ^17,18^. Additionally, we previously developed and used insertional mutagenesis through *Agrobacterium-*mediated transformation to identify regulators of filamentation and adhesion ^9,19^. However, genome-scale analyses of essentiality in *C. auris* have not yet been performed, due to difficulties in achieving saturating mutagenesis and mapping insertion sites at scale. Combining essentiality analysis with pangenomics would allow for further characterization of genes that are both conserved and essential and thus ideal as future drug development targets.

Here, we combine pangenomic analysis with global essentiality screening to build a foundational picture of the *C. auris* genome. We performed *de novo* genome assemblies for a genetically representative set of 695 genomes primarily drawn from outbreak isolates in clades I, III, and IV and used them to define the pangenome. Using ancestral reconstruction, we identified accessory gene families with clear signatures of convergent evolution, including the repeated loss of a subtelomeric deletion that overlaps a known adhesin gene. This analysis provides a detailed description of accessory gene families at both the species and clade level, in addition to offering insight into the selective pressures experienced by this population. As expected from a species that appears to have undergone a limited number of transitions to humans, we found a very high degree of clonality in our dataset of isolates, with only 2.6% of genes exhibiting clear and verifiable variation in at least 5% of the isolate genomes. Given that the vast majority of gene families were present in nearly our entire dataset, we could not infer gene essentiality from the pangenome alone. Therefore, to define the *C. auris* conserved and essential genes, we took advantage of the discovery that *C. auris* will promiscuously insert linear DNA into its genome to develop an insertional mutagenesis library and sequencing approach. We identified 614 high confidence predicted essential genes, with nearly a third showing a divergent pattern of essentiality from the model budding yeasts *Candida albicans* and *Saccharomyces cerevisiae*. As a case study, we demonstrate variability in requirement for the Sui1 translation initiation factor between *C. auris* and *C. albicans.* Together, this study provides a comprehensive characterization of the genetic variation that exists within *C. auris* through multiple parallel approaches, building a foundation for future work on identifying novel targets for antifungal therapy or species-specific determinants of virulence.

## Results

### Generating a high-confidence *C. auris* pangenome

To provide a comprehensive view of the currently sequenced *Candidozyma auris* strains, we first selected a panel of 695 genetically diverse and high-quality isolates primarily drawn from the NCBI Pathogens Detection database, which we supplemented with isolates collected from nearby hospitals (**Supplemental Table S1**). All selected isolates underwent *de novo* assembly and annotation using a custom-built snakemake pipeline^20^ designed for *C. auris* (**Supplemental Figure 1A**). We also included 41 hybrid assemblies, resulting in multiple chromosome-level assemblies for each of the three clades.

Given that *C. auris* exhibits several features more commonly associated with bacterial populations, such as a lack of sexual reproduction and highly clonal outbreaks, we performed our initial analysis with the bacterial pangenome tool Panaroo^21^, defining core gene families as those that appear in at least 95% of isolates. This resulted in 6183 gene families, 89.4% of which were core genes, as well as a network graph visualizing differences in synteny and gene order across isolates (**Supplemental Figure 1B**). Comparing these results to the common eukaryotic pangenome tool Orthofinder^22^ revealed that Panaroo reduced the total size of the pangenome by nearly 35%, while assigning more core gene families in total (5527 vs. 5370) **(Figure 1A)**. The majority of this difference was attributable to the number of singleton and near-singleton gene families, which was higher in Orthofinder by a factor of over 10X (**Figure 1A**) and was therefore less consistent with our expectations for a highly clonal population. However, consistent with previous publications^23^, we found that many accessory gene families reported by Panaroo were impacted by annotation errors. In 16-30% of clade-specific accessory gene families, every isolate where the gene was reported as absent carried the exact nucleotide sequence on the expected chromosome with no gaps or mismatches, but this region was not annotated as the expected gene. To address this annotation shortcoming, we designed a pipeline to validate gene presence and absence data using BLAST+ ^24^. Applying this tool to our dataset revealed that spurious accessory gene families were highly pervasive (**Supplemental Figure 1C**). This made it difficult to distinguish true gene loss events caused by SNPs or small indels from spurious gene loss events caused by differences in gene annotation. Therefore, we focused our downstream analysis on the subset of accessory genes where we could verify that the sequence had been lost from one or more isolates, indicating that they were caused by deletions or structural variants. After removing any accessory genes that failed BLAST validation, this resulted in a total of 144 non-core, high-confidence gene families (**Figure 1B-C, Supplemental Data 1,2**).

**Figure 1:**
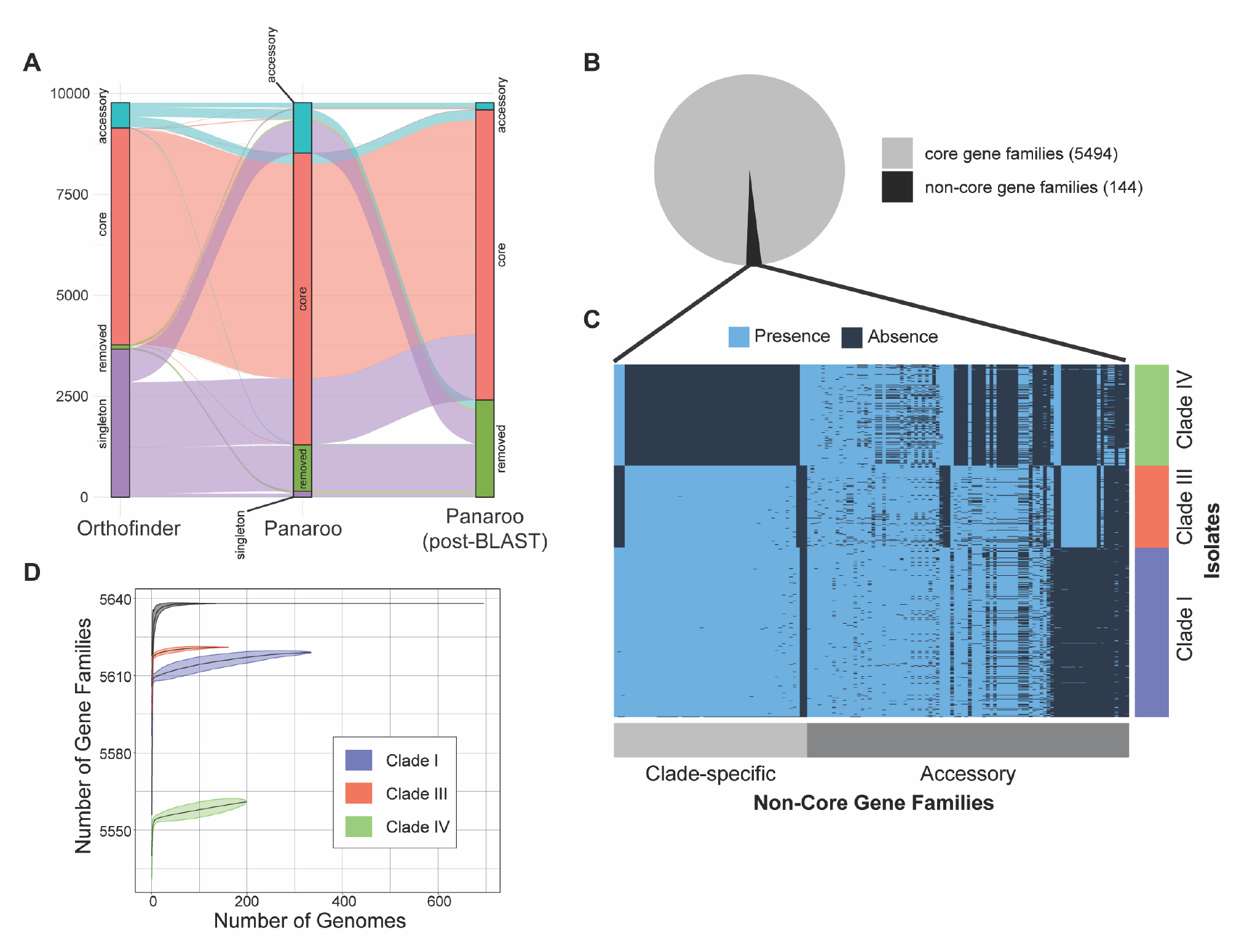
Refining the *C. auris* pangenome. **A,** Alluvial plot displaying the difference in gene family category between pangenomes produced by Orthofinder, Panaroo before BLAST validation, and Panaroo after BLAST validation. All tools were run on the same dataset of 694 *C. auris* assemblies. “Removed” indicates the corresponding gene family was either not found in the pangenome or removed via the BLAST validation step. **B,** Number of core and non-core gene families present in the pangenome after BLAST validation was performed. **C,** *C. auris* pangenome after all validation steps were performed. Only non-core gene families are shown across clades I, III, and IV. Gene families are ordered by decreasing frequency from right to left. Isolates are ordered by clade on the y-axis. **D,** Accumulation curves for the post-validation pangenome, showing the increasing size of the pangenome as more isolates are added. Clade-specific curves are shown in addition to the species-wide curve. Curves were generated via 500 random permutations, with each line representing the mean and the shaded regions indicating the 5^th^ and 95^th^ percentiles.

Following these validation steps, accumulation curves of the pangenome at both the species and clade level showed diminishing returns, indicating that our dataset had likely sampled most of the gene families present in sequenced *C. auris* isolates (**Figure 1D**). In total, we identified 5638 gene families across our 695 isolates, with an average of 5585 families per genome (sd = 32.51). We found that 97.4% of gene families were categorized as core, appearing in at least 95% of isolates, with 78.2% of gene families appearing in 100% of isolates. We identified 90 validated accessory gene families that showed variation in at least one clade. 64 of these were unique to a single clade, with 12 were shared between exactly two clades, and 14 were shared between all three. However, we also identified 54 genes that were entirely clade-specific, appearing as core in some clades and absent in the remainder. Given that a small fraction of gene families exhibited any variation in gene presence or absence,^25^ even when we designed our dataset to maximize genetic diversity, these results are consistent with a small effective population size of healthcare-associated isolates with little evidence of crossing, as in *C. parapsilosis* ^25^. This is in contrast to yeast species where crossing is more common, such as *Aspergillus fumigatus,* where only 55-69% of the genome is core ^23,26^, or *Saccharomyces cerevisiae,* where ∼60% of the genome is core ^27^.

### The *C. auris* accessory genome shows repeated gene loss

The accessory genes present in this pangenome likely include genes and loci under selective pressure, which can cause the same gene family to be gained or lost multiple independent times. As our dataset draws heavily from clinical isolates, we hypothesized that identification of the loci and genes with signatures of convergent evolution and selection in this dataset would provide insight into the selective pressures experienced by *C. auris* in healthcare settings. To accomplish this, we first generated a maximum-likelihood phylogeny tree with IQTREE ^28^, using only the single-copy core genes from the pangenome as input. We then subset our validated accessory genes based on the clade where they were identified (**Figure 2A-C**, top panels). Many accessory gene families appeared on the same chromosome and exhibited strong linkage with each other, with identical patterns of presence and absence. Of the 90 clade-level accessory genes, 46 could be grouped into clusters of two or more genes, with 15 clusters in total, resulting in 59 independent genetic loci. We then performed ancestral reconstruction to identify the number of times each locus was gained or lost (**Supplemental Figure 2A**). Roughly 18% of loci exhibited clustering on the tree, with the gain or loss events clustering together in specific subgroups (**Supplemental Figure 2A-B**), consistent with onward transmission of strains with these loss events. Contrary to our expectations, especially as we are focusing on structural variants as drivers of gene loss, several loci with high numbers of transitions were distant from subtelomeric regions (**Figure 2A-C**, bottom panels), which are known to be more prone to genomic rearrangements and deletions ^5,29,30^. However, the largest clusters of 4 or more linked gene families did all occur in subtelomeric regions, with the deletion of chromosome ends being responsible for the shared loss of several adjacent genes.

**Figure 2:**
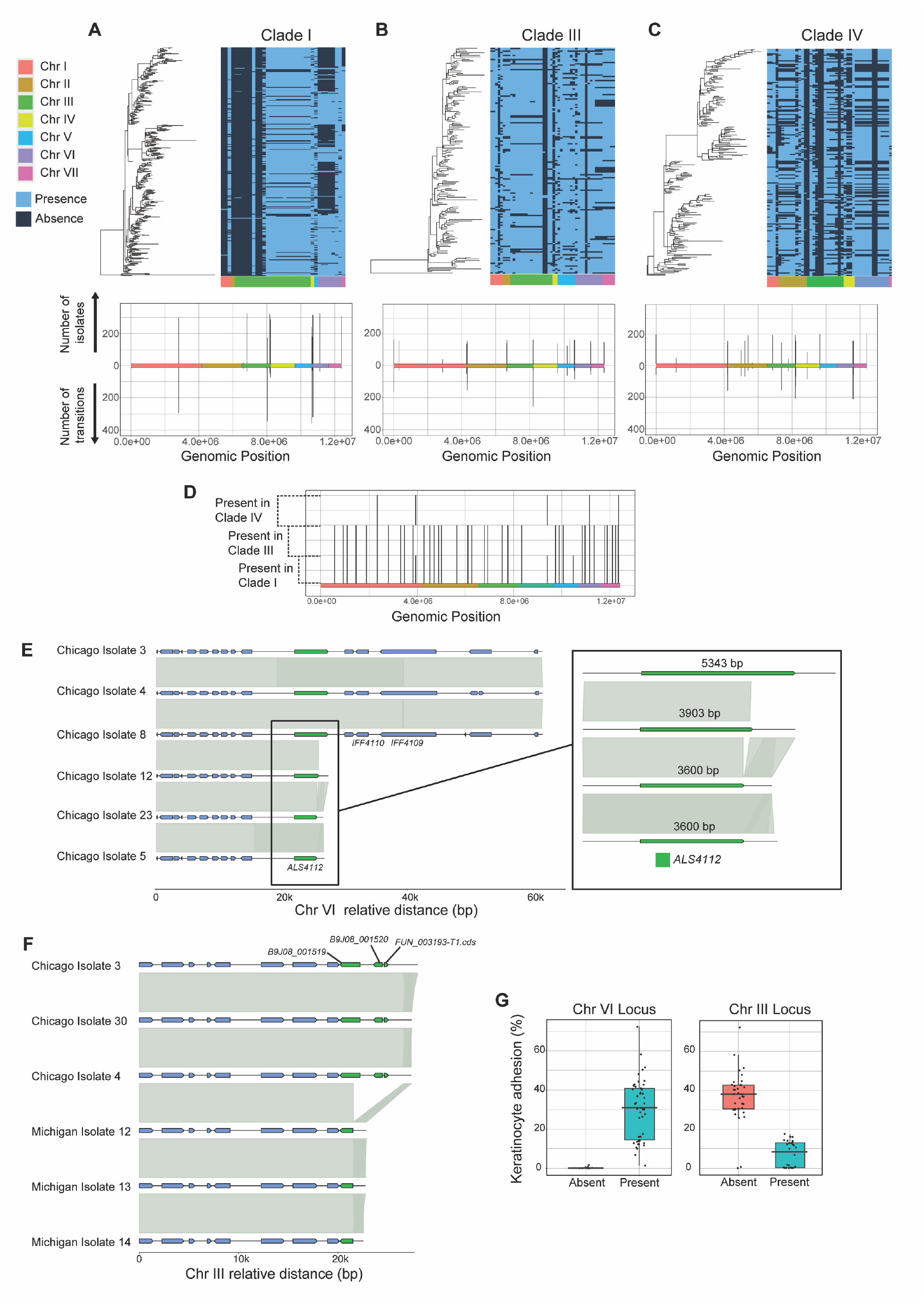
Characterization of the *C. auris* accessory genome. **A,** Top panel, left: Maximum-likelihood phylogeny tree for Clade I isolates. Top panel, right: Presence/absence matrix for all non-core gene families that are accessory in Clade I, appearing in less than 95% of isolates. Gene families are ordered on the x-axis by estimated position, based on synteny information from the pangenome. Bottom panel: Genomic location, number of isolates, and number of transitions for the accessory genes shown on the top panel. Number of transitions were determined via ancestral state reconstruction of the gene presence/absence matrix. Gene families exhibiting strong linkage are clustered together and treated as a single locus. **B,** Similar to panel A, with isolates and gene families from Clade III. **C,** Similar to panel A, with isolates and gene families from Clade IV. **D,** Estimated locations of clade-specific genes. These gene families are core in one or more clades and absent from the remainder. **E,** Representative isolates showing two different truncations of the *ALS4112* adhesin gene. All isolates shown were drawn from the Rush University Medical Center dataset. **F,** Representative isolates showing deletions on chromosome 3 associated with increased adherence. All isolates shown were drawn either from the Rush University Medical Center dataset (“Chicago”) or from the University of Michigan hospital dataset (“Michigan”). **G,** Boxplots showing the differences in keratinocyte adhesion among a subset of phenotyped clade IV isolates. Left: Isolates that either lost or retain the Chromosome VI locus that overlaps ALS4112. Presence or absence of gene family group_4112 was used as a representative for this group of tightly linked loci. Right: Isolates that either lost or retain the Chromosome III locus that overlaps three genes of unknown function. Presence or absence of gene family group_5458 was used as a representative for this group of tightly linked loci.

We next extended our analysis to gene families that did not appear as accessory in any of the three clades, but still exhibited variation at the species level. This is possible when a gene appears in nearly every member of one or more clades and is entirely absent from the others. We found a total of 54 gene families that fell into this category. Clustering of linked genes reduced this number to 41 independent loci, with the largest cluster containing 3 linked gene families. Estimating the location of these loci revealed a nearly even distribution of genes across the genome (**Figure 2G**) and did not show enrichment at chromosome ends. The majority of these loci appeared in both Clade I and Clade III, while being absent in Clade IV, consistent with the known evolutionary history of these clades ^14^.

Gene annotation data is limited in *C. auris*, but features such as InterProScan ^32^ and PFAM ^33^ terms can still provide insight into the putative functions and pathways in these clade-specific accessoriez. We found significant enrichment for InterProScan terms related to cell wall proteins (IPR021031, IPR031573), major facilitator superfamily transporters (IPR036259, IPR003663, IPR005828, IPR005829, IPR020846, IPR050360), and PIK1 kinase (IPR021601, IPR049160) among the accessory genes compared to the core genes (Bonferroni-corrected p-values < 0.05, only annotated genes tested). Significant PFAM terms across this same set of accessory genes were related to these functions, including PF11765 and PF15789 for hyphally-regulated cell wall proteins, PF00083 for sugar and other transporters, PF07690 for the major facilitator superfamily, and PF11522 and PF21245 for PIK1.

### Specific gene loss events are associated with keratinocyte attachment phenotypes

Further examination of these large clusters of linked gene families in Clade IV revealed a large locus near the end of chromosome VI that was lost multiple independent times (**Figure 2E**). Loss of this locus invariably caused truncations of the known adhesin gene, *ALS4112*, which is known to influence the clinically relevant trait of skin adhesion^9^, as well as loss of two additional adhesins *IFF4109* and *IFF4110*^8^. Notably, this deletion only appeared in isolates collected near Chicago by the Rush University Medical Center, suggesting that this locus and phenotype may be under active selection in that environment and in specific clinical contexts.

To determine the phenotypic impact of this deletion, we performed keratinocyte adhesion assays on 92 clade IV isolates from the pangenome, followed by a microbial GWAS scan using Hogwash^31^ (**Supplemental Figure 2C**). Consistent with previous studies, this revealed that the truncation of *ALS4112* via the chromosome VI deletion was significantly associated with reduced adhesion^9^ (**Figure 2G**). We also identified a second locus on chromosome III that was lost from a different subset of clinical isolates that was associated with a significant increase in adhesion when deleted, although the genes here are currently uncharacterized (**Figure 2F-G**).

### Comparing *C. auris* to the *C. haemuli* species complex

While these analyses provide detailed insight into the genetic variation that exists in the accessory genome of *C. auris*, the highly clonal nature of our dataset means that we cannot assume that core gene families are maintained due to the essentiality of their biological functions. It is likely that the core gene families appearing in the 95% or more of our isolates consist of a combination of essential and nonessential genes, given how infrequently we observe presence and absence variation in our pangenome, even before BLAST validation. This is supported by the fact that only 43.9% of the core genome from *C. auris* is present in at least two related species among *C. haemuli* and its nearest relatives (**Figure 3**). To further characterize and differentiate this large category of putative core gene families, we used the complementary approach of essentiality mapping via insertional mutagenesis.

**Figure 3:**
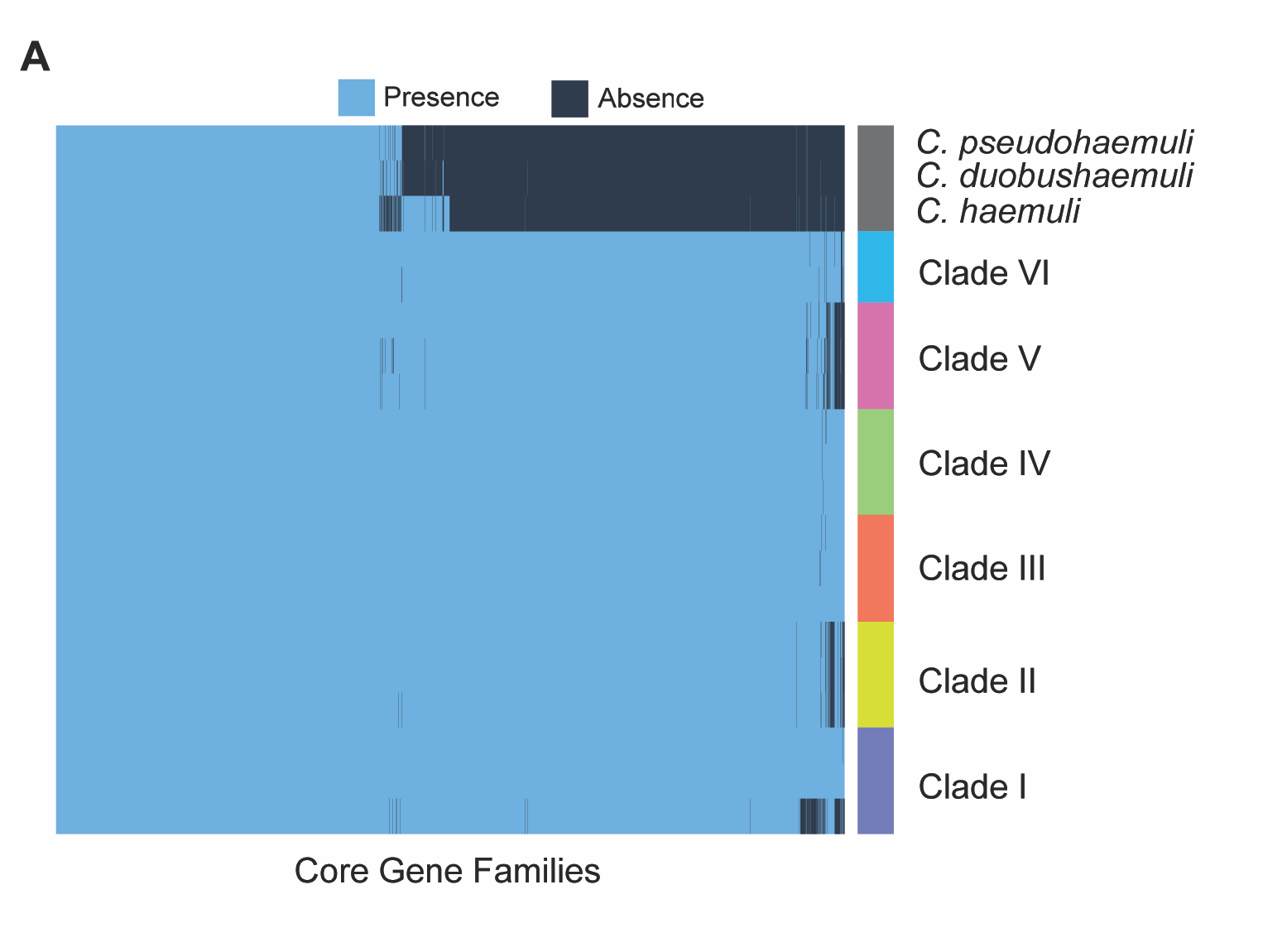
*C. auris* core genome compared to *C. haemuli* species complex. Results of a pangenome generated using 703 genomes from all six clades of *C. auris*, as well as reference genomes for *Candida haemuli*, *Candida duohaemuli*, and *Candida pseudohaemuli*. For each clade, 2-3 representative isolates are shown, prioritizing the highest-quality assemblies when possible. Only gene families categorized as core (present in >95% of isolates) are shown on the heatmap, and are ordered on the x-axis based on decreasing frequency from left to right.

### Transformation of a selection marker throughout the *C. auris* genome

Precise genetic editing through transformation in *C. auris* is challenging; despite large homologous arms, homologous recombination is rare and instead the cassettes are incorporated in an off-target site instead of cleanly mutating a target gene ^19,34^. We leveraged this off-target integration to perform insertional mutagenesis by transforming a nourseothricin (NAT) resistance cassette throughout the Clade I isolate AR0382 genome and using transposon footprinting to map insertions (**Supplemental Figure 3**). From fifteen individual libraries, we identified a total of 208,333 unique NAT cassette insertion sites widely distributed across all seven *C. auris* chromosomes (**Figure 4A**). Similar to other transposon-based studies, we observed a strong bias towards repetitive regions, including rDNA repeats. We account for this by focusing our analysis on genic NAT cassette inserts ^35^. These NAT inserts did not display a detectable bias towards any sequence motif enrichment at the sequenced NAT 3’ end. (**Figure 4B**). To examine insertion bias by chromatin accessibility, we performed ATAC-seq and observed enrichment at accessible sites (**Figure 4C**). Additionally, whole genome sequencing of individual colonies also identified only a single insertion event per genome. Holistically, this supports that our method is comparable to traditional transposon mutagenesis strategies, and defines a novel method for *C. auris* insertional mutagenesis.

**Figure 4:**
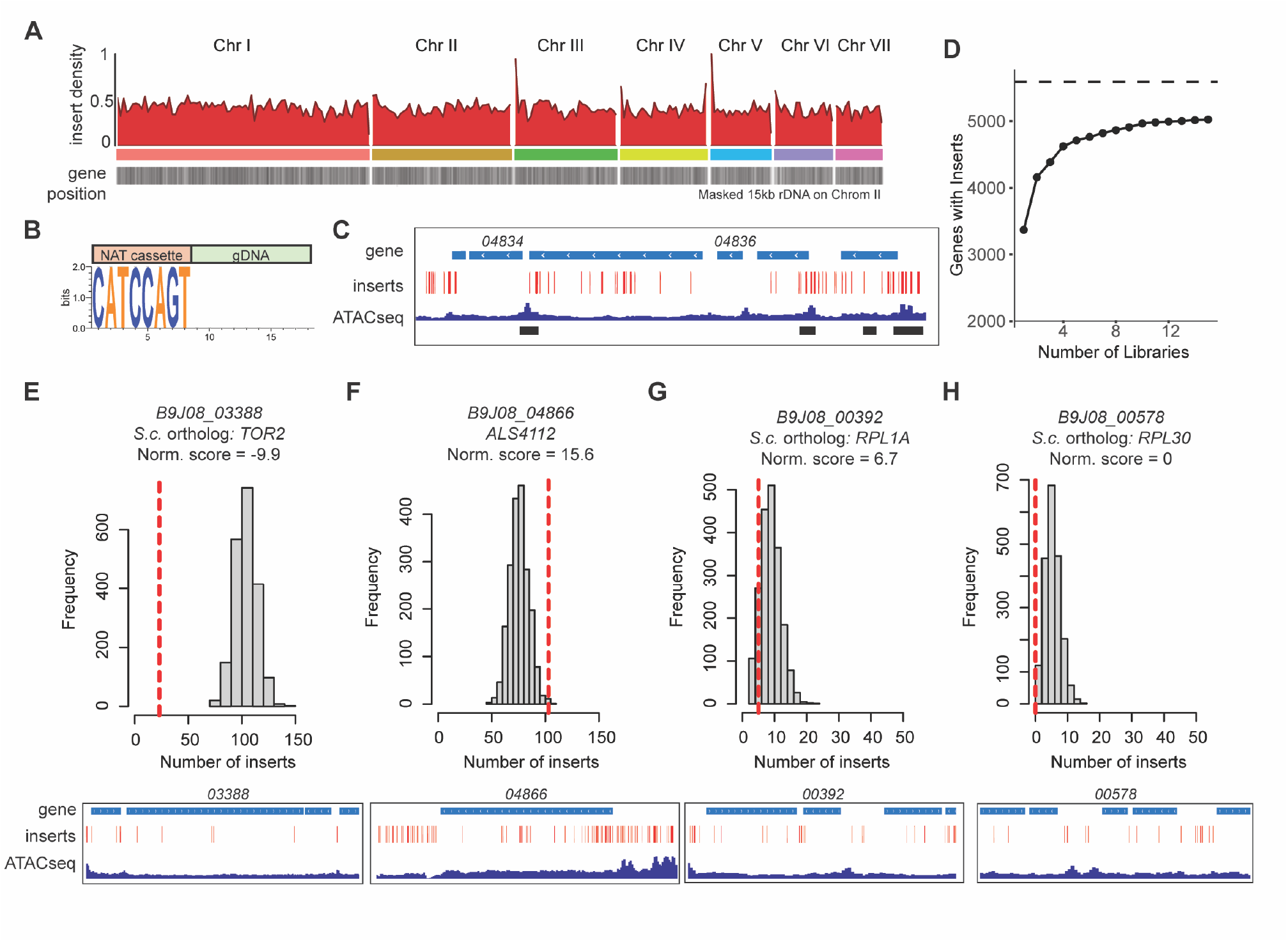
Transformation of a selection marker throughout the *C. auris* genome. **A,** Karyoplot illustrating NAT-resistance gene insertion density (red) per 50kb-bin across the *C. auris* genome at over 200k unique insertion sites. Gene density is shown below (gray). 15kb on chromosome CM076439.1 was masked to improve visible scale. **B,** Sequence logo generated from the hybrid NAT-gDNA insertions. ‘CATCCAGT’ corresponds to the end of the NAT resistance cassette. **C,** IGV tracks displaying select *C. auris* genes at the indicated coordinates (top). Insert sites, ATAC-seq, and MACS3-identified ATAC-seq peaks are shown below. **D,** Number of gene ORFs containing at least one NAT insert site reached an asymptote with additional libraries. **(E,F,G,H,)** (top) Random permutation simulation analysis distributed the proportion of inserts landing in coding ORFs (∼115k sites) across the gene sample space. Comparing the lower bound of the distribution to our observation (red line) allowed us to further identify ORFs of essential genes. (below) IGV tracks displaying select *C. auris* genes, insert sites, chromatin accessibility (ATACseq), and transcription (RNAseq) at the indicated coordinates.

### Essentiality mapping in *C. auris*

As we increased our number of insertional libraries, we observed a saturation for the number of genes tolerating interruptions, suggesting insertion-deplete regions may be essential (**Figure 4D**). For example, despite accessibility at *B9J08_04834* and *B9J08_04836*, these genes are insert-poor, consistent with their conserved essential roles in mRNA 5’ guanylyltransferase capping and the U2 snRNP complex, respectively ^36^. However, the lack of a defined insertion motif precluded us from using existing transposon mutagenesis essentiality methods ^37^. Additionally, other work using transposon mutagenesis methods have found that some essential genes can tolerate interruptions ^16,35^. Therefore, to conservatively predict gene essentiality, we simulated repeated random redistribution of genic NAT inserts (∼115k) across the *C. auris* coding region and normalized the expected number of inserts to the gene length. To be considered essential, we set a conservative threshold of at least five inserts less than the lowest value of the permutations or zero observed inserts. As an example, under these criteria, we observe significantly fewer insert sites in the conserved essential gene *TOR2* compared to the simulated distribution (**Figure 4E**). In contrast, genes we have previously experimentally determined to be non-essential, like *ALS4112* (*B9J08_04866*), do not deviate from a random distribution of inserts (**Figure 4F, Supplemental Figure 4**) ^9,19^. As a limitation, the simulated insert distribution in short genes (< 700 bp) includes zero inserts, so we categorize these as unresolvable (UN) (**Figure 4G,H**).

### *C. auris* gene essentiality highlights divergent gene dependence

We predicted 614 essential genes and 219 as unresolvable (**Figure 5A, Supplemental Data 3**). From our predicted essential genes, we found none of them were validated accessory genes in the pangenome. Additionally, these essential genes were significantly enriched among the core genes that *C. auris* shares with *C. haemuli* and close relatives (**Figure 3A**, Fisher’s exact test p-val = 3.3 x 10^-5^).

**Figure 5:**
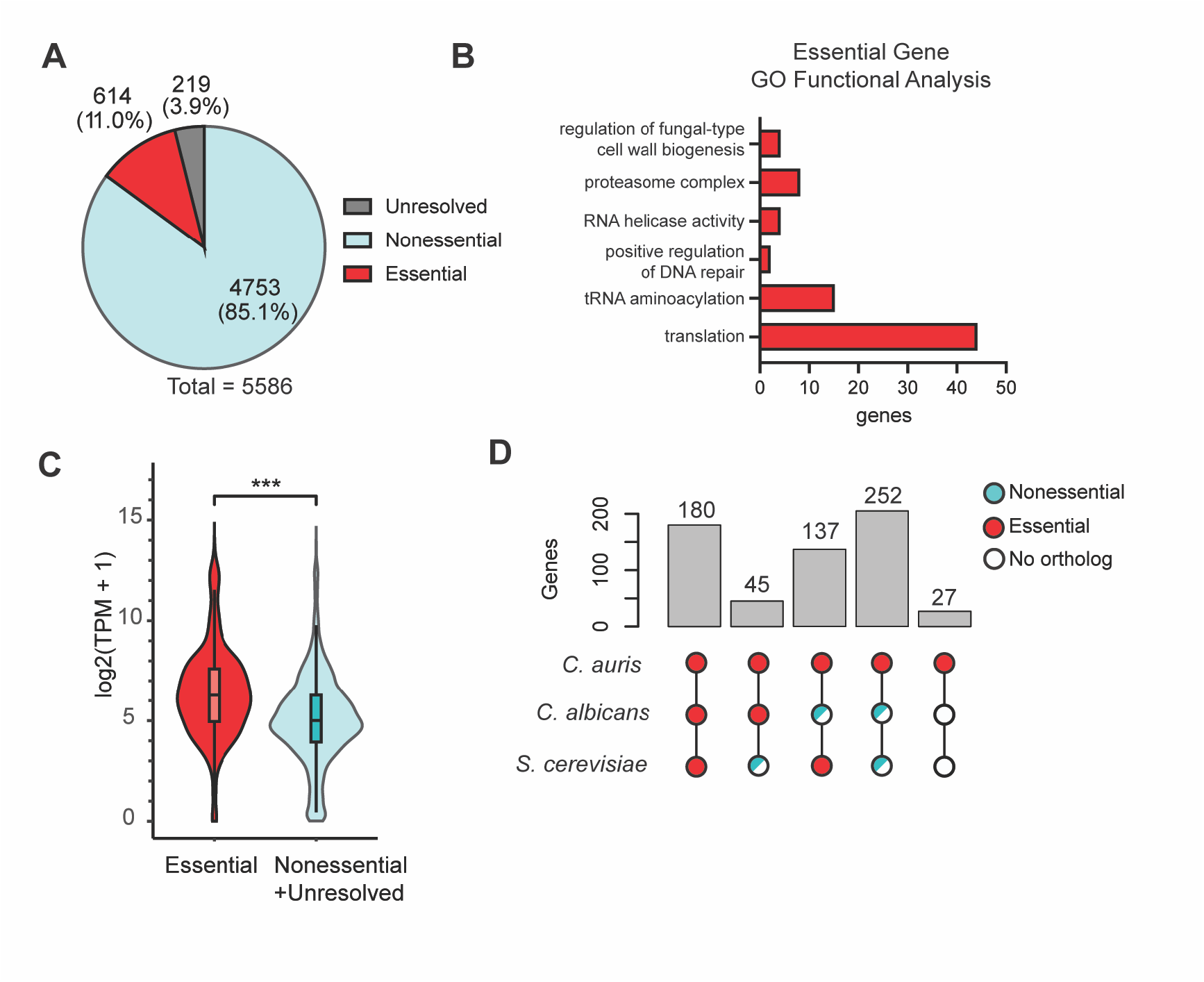
*C. auris* gene essentiality highlights divergent gene dependence. **A,** *C. auris* genes that are essential, nonessential, or unresolved by insertional mutagenesis analysis. **B,** Significantly enriched terms from GO functional analysis of 614 essential *C. auris* genes, performed using the Candida Genome Database GO Term Finder. **C,** RNAseq expression in log2(TPM+1) for *C. auris* essential and nonessential genes. Unresolved genes were binned with nonessential genes. Data from Santana et al. 2023. **D,** Comparison of 614 *C. auris* essential genes to orthologs in *C. albicans* and *S. cerevisiae*. 47 *C. auris* putative essential genes had no annotated ortholog to either related species, and are a subset of the previous group of 252 essential *C. auris* genes that are nonessential or have no ortholog in *C. albicans* and *S. cerevisiae*.

For essential *C. auris* genes, GO analysis highlighted enrichment for conserved essential functions (**Figure 5B**). Unresolved genes were similarly enriched for essential functions, including RNA polymerase activity, snRNA binding, and ribosome biogenesis (**Supplemental Figure 5A**). To address the lack of GO term annotation for *C. auris*, we repeated GO term analysis using available *C. albicans* orthologs, which revealed additional enrichment for actin cytoskeleton organization and phospholipid biosynthesis **(Supplemental Figure 5B).** Lastly, essential genes tend to exhibit higher expression ^38,39^. Therefore, we compared the expression levels of our predicted essential and nonessential gene sets from a previously published RNAseq dataset^8^ and observed that our predicted essentials have higher transcript levels (**Figure 5C**). Collectively, this supports our capacity to define essentiality and their correspondence with essential functions.

### *C. auris* essentiality diverges from model yeasts

Since our permutation-based analysis does not incorporate essentiality information from other species, we can perform unbiased comparisons to related fungi and identify evolutionary divergences in *C. auris* essentiality. We compared essentiality and ortholog data to *S. cerevisiae* and *C. albicans,* where extensive work has been previously performed ^36,40–43^. Overall, we found moderate agreement where 180/614 essential *C. auris* genes had essential orthologs in both *C. albicans* and *S. cerevisiae* (**Figure 5D**). We performed a chi-square test of independence to examine the conservation of essentiality across orthologs and observed a significant relationship (*χ*2(1,N=614) = 114.5, p<0.001). However, we identified a third of our putative essential genes in *C. auris* that were nonessential in *C. albicans* or *S. cerevisiae*. Though annotation of the divergently essential gene set was limited, GO term analysis revealed enrichment for cell replication and cellular respiration terms (**Supplemental Figure 5C**). This is consistent with the ability of both *C. albicans* and *S. cerevisiae* to grow anaerobically, while *C. auris* cannot ^44^. Lastly, 27 of our putative *C. auris* essential genes do not have clear sequence orthologs to our two comparison species, precluding a comparative analysis of essentiality. All 27 of these lineage-restricted genes are expressed (TPM >= 1) (**Supplemental Figure 5D**). These divergences in essentiality highlight the variable dependence on individual genes between these three fungal species.

To experimentally test a putative essential locus, we chose the DASH complex subunit *Spc19* ortholog *B9J08_02579* to put under tetOFF-repressible promotion ^45^. The fungal-specific DASH complex ensures proper chromosome segregation during closed mitosis, though its essentiality depends on the nature of kinetochore-microtubule interactions during the cell cycle ^46^. Initial growth on doxycycline-containing media displayed nearly complete growth inhibition, strongly supporting the inferred essentiality of this locus (**Figure 6A**). Through microscopy, we observed that *Spc19* repression in both *C. auris* and *C. albicans* resulted in elongated morphology, consistent with cell cycle arrest (**Figure 6B**) ^47,48^ ^49^.

**Figure 6:**
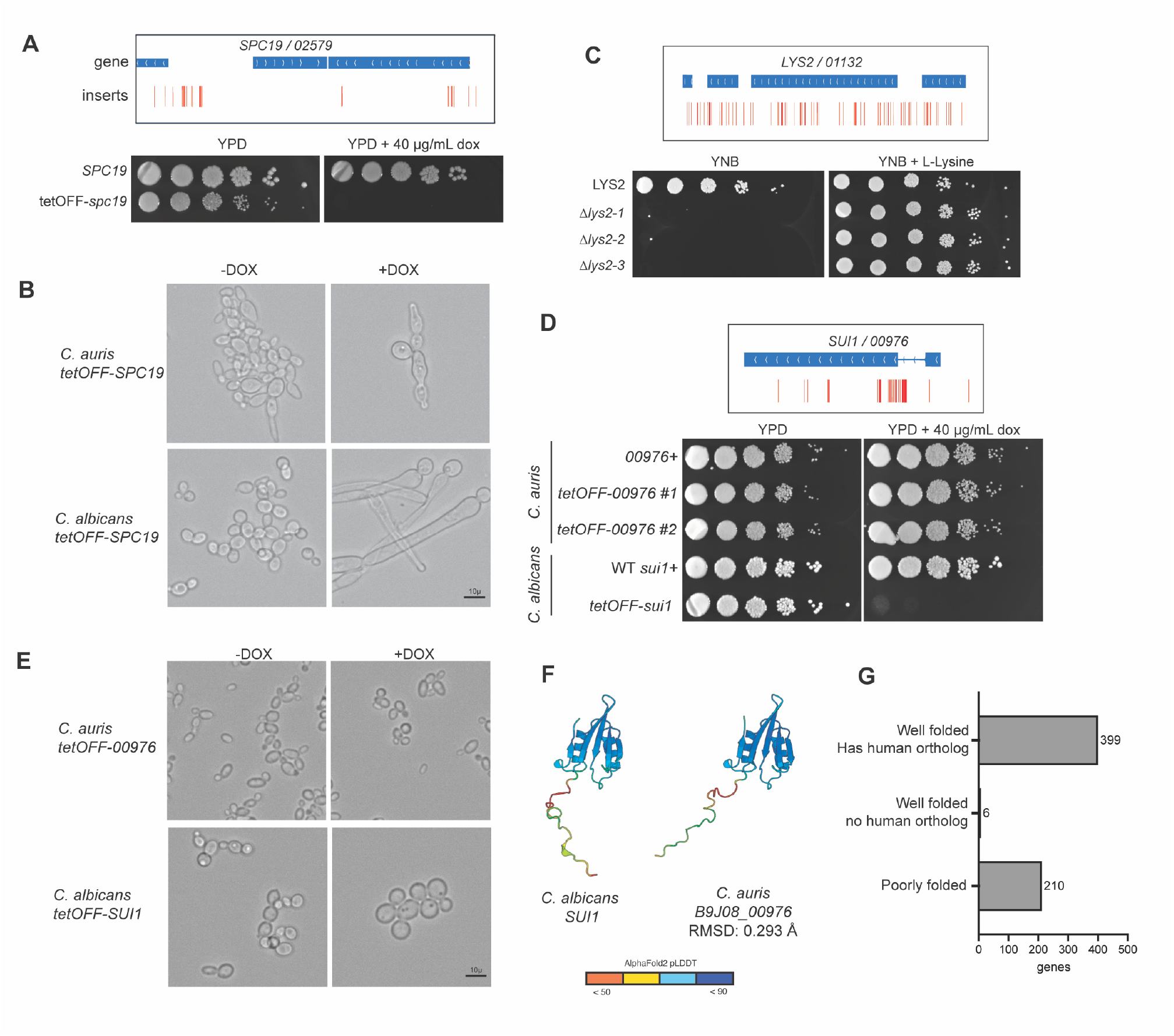
Validation of *C. auris* gene essentiality. **A,** IGV tracks for the *SPC19* locus and a spot plate showing growth for the indicated strains under a tetOFF promoter in the presence or absence of doxycycline. **B,** Morphology of the tetOFF-SPC19 strains *C. albicans* and *C. auris* in the presence or absence of doxycycline in YPD. **C,** IGV tracks at the *LYS2* locus and a YNB agar spot plate showing growth in the presence or absence of lysine. **D,** IGV tracks for the *SUI1* locus and a spot plate showing growth for the indicated strains under a tetOFF promoter in the presence or absence of doxycycline. **E,** Morphology of the indicated strains in the presence or absence of doxycycline. **F,** Alphafold-predicted structures of the *C. auris* or *C. albicans* Sui1 proteins. **G,** Structural analysis of *C. auris* essential genes identifies candidates without structural orthologs in humans.

We also identified 664 *C. auris* nonessential genes that had essential orthologs in related species. This is partially due to the conditions which essentiality was determined for each species, wherein the studies conducted for gene essentiality in *C. albicans* and *S. cerevisiae* were performed in minimal media to identify genes for nutrient synthesis ^16,41,50^. However, under rich media conditions in our method, essential nutrient synthesis genes appear nonessential. To experimentally validate this, we generated a deletion strain of the *C. auris LYS2* ortholog *B9J08_01132* (**Figure 6C**). This strain exhibits auxotrophy on minimal media, which can be rescued through the supplementation of L-Lysine. This supports the sensitivity of our mutagenesis to properly identify gene nonessentiality in rich media.

However, we also identified cases where the discordance in gene essentiality suggests an evolutionary divergence in gene dependencies. To exemplify this, we generated a repressible allele of the *SUI1 C. auris* ortholog, *B9J08_00976* ^51^. In related species, Sui1 acts as translation initiation factor eIF1, which is an essential component for eukaryotic protein synthesis ^52^. On doxycycline-containing media, we do not observe any growth defect or morphological changes in *C. auris* tetOFF-*00976*, contrasting with the swollen morphology and poor growth of the analogous tetOFF-*sui1 C. albicans* strain (**Figure 6D-E, Supplemental Figure 6A**). The mechanism that allows for dispensable *SUI1* in *C. auris* is unclear, especially as the proteins show high sequence and structural similarity to the *C. albicans* protein, and we did not identify any paralogs or duplications through BLAST that would confer redundancy **(Figure 6F)**. Notably, a protein containing an SUI1-like domain was present in *C. auris,* but the rest of the protein structure was not shared **(Supplemental Figure 6B).** Altogether, our experimental data supports a model where *C. auris* gene essentiality and translation biology has diverged from related species ^50,53^.

Lastly, identifying gene essentiality is key to future antifungal development. We performed a structural similarity search using FoldSeek to identify essential *C. auris* genes lacking structural orthologs in humans, identifying targets with low likelihood of concurrent host toxicity (**Figure 6G**)^54^. From our search, we identified 6 *C. auris* essential genes that have high confidence structures (pLDDT >= 70) and lack a structural human ortholog (TM < 0.5). Of these six, two are previously identified as possible drug targets, including *B9J08_04245*, a fungal mitochondrial subunit ortholog, and *B9J08_01210*, a tryptophan synthase ortholog. However, 209 of the *C. auris* essential proteins had low-confidence predicted structures, limiting our ability to look for structural orthologs.

## Discussion

In this study, we leverage both essentiality mapping and pangenome analysis to characterize the *Candidozyma auris* genome. Our pangenome analysis identifies high-confidence accessory gene families, many of which exhibit strong phylogenetic signals and can be grouped into clusters of linked genetic loci. However, this approach is limited by low genetic variation in our highly clonal clinical isolate dataset and sparse functional gene annotations. To complement these limitations, we took advantage of the high off-target transformation rate of *C. auris* to develop a novel gene essentiality mapping technique, providing functional context for many of these gene families.

Inflation of pangenome size through the addition of spurious accessory or singleton gene families is a known problem in pangenomic studies ^55^. Small inconsistencies in gene annotation can cause a core gene family to be incorrectly split into multiple accessory genes, even when there are no changes at the nucleotide level. However, true gene loss events can also be caused by as little as a single nucleotide change, such as in the case of premature stop codons. Rather than attempting to distinguish between these two cases, we instead chose to focus our analysis on the set of accessory genes that were caused by deletions or structural variants. These can be identified and validated at high confidence, as the expected nucleotide sequence is partially or entirely missing from one or more isolates. We found that this approach significantly improved the consistency of the pangenome with existing knowledge of *C. auris*. Validating our accessory genes in this way retained many of the gene loss events and excluded many of the gene gain events, which matches our expectations of a fungal species that does not demonstrate meiotic recombination ^56^. This consistency also extends to our own essentiality mapping, where none of the Clade I accessory genes overlap with predicted essential genes.

We initially hypothesized that the highest-confidence accessory genes would be entirely localized to subtelomeric regions, which experience genomic instability and are the easiest to verify. However, we instead found that accessory genes are distributed evenly across the genome, implying that our results cannot be explained simply by chromosomal instability in subtelomeric regions. In future studies, we aim to combine this dataset with additional types of genetic variation, such as structural variants, SNPs, and indels, to better understand the genomic events leading to the presence and absence variation we observe here.

In addition to providing characterization of the core genome, insertional mutagenesis allowed us to identify *C. auris* essential genes that lack any known orthologs in humans. As fungus-specific proteins, these provide potential targets for future studies into antifungal drug development. We identified 6 essential genes with high structural quality scores and no structural orthologs in humans, including *B9J08_04245*/*Nam9,* a mitochondrial small ribosome subunit, and *B9J08_01210/Trp5,* a tryptophan synthase. The mitoribosome has been recognized as a favorable therapeutic co-target, where mitoribosomal inhibition confers fluconazole hypersensitivity in *C. albicans* ^57^. Similarly, tryptophan synthase has been considered as an antimicrobial target due to its lack of human homologs ^58–60^. Importantly, 209 proteins had poor quality structure prediction, potentially due to a lack of training data for lineage-specific proteins in the AlphaFold models. Nevertheless, this highlights additional putative essential gene targets for antifungal development.

Promiscuous gene incorporation generates mutant libraries without exogenous genetic toolkits. However, unlike transposon-based mutagenesis that target-specific microsequences, it is unclear whether *C. auris* promiscuous gene insertion is sequence-dependent ^61^. Since we do not have prior knowledge of the insertion space, we are restricted from performing typical transposon distribution analyses through Bayesian methods, which necessitates a more conservative analysis ^62^. In addition, we outgrow the mutant pools in a competitive liquid growth environment, which may select against genes whose deletions produce defects in planktonic growth i.e. excessive flocculation.

Our insertional mutagenesis method implies that *C. auris* can readily integrate foreign DNA into its genome. However, our pangenome indicates that this is not a major driver of the genetic variation observed in our dataset of clinical isolates. Despite focusing our analysis almost entirely on accessory genes caused by structural variants, the majority of our data was consistent with gene loss rather than gene gain. Further work is required to determine whether this integration potentially highlights a novel pathway for generating genetic variance in *C. auris* ^56^.

## Supporting information

sup data 1 pangenome

sup data 3 essential genes

sup data 2 pangenome cds names

sup data 2 pangenome names zipped

## Acknowledgements

We thank all members of the Snitkin and O’Meara labs for helpful comments. We thank MDHHS, Mary Hayden, and the UM Infection Prevention team for sharing *C. auris* strains. Funding for this work was provided by:

AJL: MMMP T32AI007528, Michigan Postdoctoral Pioneer

TRO and ES: U19AI181767

TRO: R35GM147894

TRO: OORRS033123 University of Michigan

JRR: T32GM007544

GZ: F32AI181164, Michigan Postdoctoral Pioneer

LF: NIH R35GM128637, NIH R01AI134678

## Methods

### Public Data and Sequencing

All public data used in this study was downloaded from the NCBI Pathogens database. Published SNP trees were used to select isolates from this database that differed by at least 30 SNPs. When possible, at least one isolate was selected from every SNP tree. All isolates not obtained from NCBI were sourced from either Rush University Medical Center or University of Michigan Medicine. Isolates from these sources were sequenced via Plasmidsaurus yeast hybrid sequencing orders, with library preparation performed as described below (section ‘Library preparation and sequencing’).

### Assembly and Annotation

All genomes used in this study were assembled and annotated via the funQCD snakemake pipeline (https://github.com/Snitkin-Lab-Umich/funQCD). A detailed description of the steps and tools used is available on the GitHub repository. To briefly summarize, raw reads were cleaned via Trimmomatic and assembled via Spades. Assembled genomes were used as input for the fungal genome annotation tool Funannotate ^26^. RNA-seq data from a panel of isolates was used as training data for structural annotation steps. GeneMark ^63^, InterProScan ^32^, and EggNOG-mapper ^64^ were used to improve functional annotations. These functional annotations are added coding sequences during the *de novo* assembly pipeline, and use protein homology to infer possible functions. Auriclass v0.5.4 (https://github.com/RIVM-bioinformatics/auriclass) was used to determine the clade of the isolate, and BUSCO ^65^ was used to evaluate completeness of both the annotation and assembly steps. MultiQC was used to aggregate QC data for downstream data cleaning.

### Initial Pangenome Analysis

Pangenome analysis was performed with both Orthofinder ^22^ and Panaroo ^21^. Protein .fasta files produced as output by Funannotate during the funQCD workflow were used as input for Orthofinder. Nucleotide .fasta and annotation .gff files from the same pipeline were used as input for Panaroo. As Panaroo was originally designed for bacterial genomes, it cannot process the ∼5% of *C. auris* genes that contain introns. To account for this issue, we spliced introns out of these genes before providing input files to Panaroo. In the rare cases where introns could not be spliced due to overlapping another gene (<50 genes across all assemblies, <0.001%), the intron-containing gene was removed from analysis. Comparing Panaroo output before and after this splicing process verified that ∼5% more genes per isolate were recovered, without significantly affecting the remaining gene families. Both Orthofinder and Panaroo were otherwise run with minimal changes from default settings.

### BLAST Validation

All accessory and clade-specific gene families were validated via the pangenome_blast_smk pipeline (https://github.com/Snitkin-Lab-Umich/pangenome_blast_smk), primarily built around the command-line BLAST+ tool. A detailed description of the steps and functionality of this pipeline is available on GitHub. To briefly summarize, a custom BLAST database was first generated for all isolates. For each gene family, a single query sequence is extracted from an isolate that carries that gene, with priority given to high-quality hybrid assemblies. This query is BLASTed against the entire subject database, regardless of which isolates the gene originally appeared in the pangenome. Only hits with a sequence identity over 90%, query coverage over 90%, and e-values lower than 1e-5 are considered valid. In parallel, the pangenome graph from Panaroo is used to determine the closest neighbors of the target gene in each isolate, using synteny information. If within-clade accessory genes are being validated, BLAST hits are additionally required to appear on the same chromosome or contig within 50 kb of these expected neighboring genes. Species-wide searches do not use this threshold, as we observed less accurate synteny and neighbor information when performing cross-clade comparisons. BLAST hits are separated based on whether the gene family was originally reported as a presence or an absence in the pangenome for each isolate. Any gene families where more than 20% of the reported gene absences are found as valid hits via BLAST are considered spurious. As an additional requirement, at least one of the reported gene presences must be consistent with BLAST. Agreement between BLAST hits and pangenome presence data was used as a positive control, with roughly 2% of gene families displaying significant inconsistencies, indicating that the pipeline was performing accurately and as expected in this dataset.

### Gene Family Location Estimation

Many gene families do not appear in reference genomes, even when using clade-specific references. For these gene families, the pangenome graph from Panaroo was used to infer its approximate genomic location. This was done using the find_pangenome_neighbors.py script within pangenome_blast_smk, as described above. To summarize, locations were determined by starting at the node representing the gene family of interest and traversing through the graph until a gene family was found that did appear in the reference genome. This process was repeated up to two times, depending on the number of edges connected to the node. If two neighbors were found, the approximate location of the target gene family was set as the interval between the two neighboring genes. If only one neighbor was found, the approximate location was set as the location of the only neighbor.

### Strains and Culture Conditions

All strains used in this study are listed in **Supplemental Table S2**. Cells were cultured at 30°C in YPD or YNB base media, and liquid cultures were constantly agitated during incubation. When required, media was supplemented with 200 μg/mL nourseothricin, 40 μg/mL doxycycline, or 50 μg/mL L-Lysine. All strains were maintained as frozen stocks in 25% glycerol at 80°C. Microscopy was performed on a BioTek Lionheart FX Automated Microscope.

### Primers and Plasmids

Supplemental tables contain lists of primers (**Supplemental Table S3**) and plasmids (**Supplemental Table S4**) used in this study. Plasmids constructed from fragments were assembled using the NeBuilder HIFI DNA Assembly Master Mix (NEB #E2621) according to the manufacturer manual.

### *C. auris* Transformation

*C. auris* transformation was performed as described previously ^19^. Briefly, PCR was used to amplify linear cassettes of the repair, the targeting sgRNA, and Cas9. Each piece was purified using a Zymo DNA Clean & Concentrator kit (Cat no. D4034, Zymo Research) according to the manufacturer’s instructions.

To prepare electro-competent cells, *C. auris* cells were grown overnight in YPD at 30 °C, pelleted, resuspended in TE buffer with 100 mM Lithium Acetate and incubated at 30 °C for 1 hr with constant shaking. Then, DTT was added at a final concentration of 25 mM and the cells were incubated at 30 °C for 30 min. Cells were harvested by centrifugation at 4 °C before being washed once with ice-cold water and once with ice-cold 1 M Sorbitol. Harvested cells were resuspended in ice-cold 1M sorbitol and kept on ice for transformation

For electroporation, 45 μL of competent cells were added to a pre-chilled 2 mm-gap electro-cuvette (Biorad) along with 500-1000 ng each of the linear cleaned DNA, and then cells were electroporated using a Bio-Rad MicroPulser Electroporator according to the pre-defined *P. pastoris* (PIC) protocol (2.0 kV, 1 pulse). The cells were then immediately recovered in 1 M cold Sorbitol, then moved to YPD and allowed to outgrow before selecting.

### Strain Construction

*C. auris* mutant strains were generated using a transient CRISPR system ^19^. The universal Cas9 cassette was amplified from plasmid pTO135 using primers oTO143 and oTO41. Plasmids with repair cassettes were assembled using the NEBuilder HIFI DNA Assembly Master Mix (NEB #E2621) according to the manufacturer’s instructions. Specific mutant colonies were identified through colony PCR using Phire Plant Direct PCR Master Mix (F160; Thermo Fisher Scientific) according to the manufacturer’s instructions. Each mutant was confirmed by at least three independent PCR reactions with distinct primer sets specific to the mutation site and compared to parental strains.

#### pTO505/tetO-B9J08_00976

To generate t*etO*-*B9J08_00976* strains, the gRNA cassette was constructed by Splice-On-Extension PCR, incorporating a 20-bp guide sequence targeting the endogenous promoter (-203∼-1) of *B9J08_00976* via primers oTO3255 and oTO3256. To generate plasmid with the repair template for t*etO*-*B9J08_00976*, the plasmid backbone was amplified from pTO139 using primers oTO590 and oTO591. The 500 bp upstream sequence (-703∼-204) of *B9J08_00976* and the 500 bp gene sequence (+1∼+500) were amplified from AR0382 genomic DNA using primer sets oTO3249/3250 and oTO3251/3252, respectively. The *tetO* cassette with a NAT selective marker was amplified from pTO240 with primers oTO3124 and oTO3125. The four fragments were assembled to generate a plasmid with the repair template for t*etO*-*B9J08_00976.* The repair cassette was amplified from pTO505 using primers oTO3259 and oTO3260 and transformed into AR0382 strain to generate t*etO*-*B9J08_00976* strains.

#### pTO487/tetO-B9J08_02579

To generate t*etO*-*B9J08_02579* strains, the gRNA cassette was constructed by Splice-On-Extension PCR, incorporating a 20-bp guide sequence targeting the endogenous promoter (-223∼-1) of *B9J08_02579* via primers oTO3074 and oTO3075. To generate plasmid with the repair template for t*etO*-*B9J08_02579*, the plasmid backbone was amplified from pTO139 using primers oTO590 and oTO591. The 500 bp upstream sequence (-735∼-224) of *B9J08_02579* and the 500 bp gene sequence (+1∼+501) were amplified from AR0382 genomic DNA using primer sets oTO3068/3069 and oTO3070/3071, respectively. The *tetO* cassette with a NAT selective marker was amplified from pTO240 with primers oTO3124 and oTO3125. The four fragments were assembled to generate a plasmid with the repair template for t*etO*-*B9J08_02579.* The repair cassette was amplified from pTO505 using primers oTO18 and oTO19 and transformed into AR0382 strain to generate t*etO*-*B9J08_02579* strains.

#### pTO540/tetO-B9J08_01132

To generate *B9J08_01132::NAT* strains, the gRNA cassette was constructed by Splice-On-Extension PCR, incorporating a 20-bp guide sequence targeting the CDS of *B9J08_01132* via primers oTO3098 and oTO3099. To generate plasmid with the repair template for *B9J08_01132::NAT*, the plasmid backbone was amplified from pTO139 using primers oTO590 and oTO591. The 500 bp upstream sequence (-1∼-500) of *B9J08_01132* and the 500 bp gene sequence (+4177∼+4677) were amplified from AR0382 genomic DNA using primer sets oTO3092/3093 and oTO3094/3095, respectively. The NAT selective marker was amplified from pTO128 with primers oTO668 and oTO1093. The four fragments were assembled to generate a plasmid with the repair template for *B9J08_01132::NAT.* The repair cassette was amplified from pTO540 using primers oTO18 and oTO19 and transformed into AR0382 strain to generate *B9J08_01132::NAT* strains.

### Genomic DNA Isolation

Genomic DNA was isolated as previously described ^8^, using a phenol chloroform extraction method. Overnight cultures were harvested by centrifugation and resuspended in lysis buffer (2% (v/v) Triton X-100, 1% (w/v) SDS, 100 mM NaCl, 10 mM Tris-Cl, 1 mM EDTA). Cells were disrupted by bead-beating and DNA was extracted using phenol chloroform, purified through ethanol precipitation and treated with RNase A (Qiagen, cat no. 19101). Rnase was heat-inactivated, then extracted DNA was purified by ethanol precipitation and resuspended in water.

### RNA extraction and RT-qPCR

RNA extraction and qRT-PCR was performed using a formamide extraction method, as previously described ^8^. Cells were incubated with or without 40 ug/mL DOX overnight in YPD, harvested by centrifugation, and snap frozen at -80 °C before processing. For RNA extraction, pellets were resuspended in 100 μL FE Buffer (98% formamide, 0.01 M EDTA). 50 μL of 500 μm RNAse-free glass beads was added to this suspension and the mixture was homogenized for 30 sec 3 times using a BioSpec Mini-Beadbeater-16 (Biospec Products Inc., Bartlesville, OK, USA). The lysate was then clarified by centrifugation and RNA was purified using the Qiagen RNeasy mini kit (ref 74104, Qiagen) according to the manufacturer’s instructions. Samples were DNAse treated with Qiagen DNAse (Qiagen, cat no. 79254).

### ATACseq

To prepare spheroplasts, *C. auris* grown overnight in liquid culture was diluted and grown out for six hours to allow return to logarithmic growth. Approximately 1.6 OD of cells were harvested through centrifugation at 13,000 rpm for 1 minute. Supernatant was aspirated, and the pellet was washed in 1 mL ice cold water then centrifugation repeated at 13,000 rpm for one minute. Water was aspirated, pellet was resuspended in buffer 1 (100mM Tris pH 9.4, 10mM DTT) and incubated for 5 minutes at 30°C. Sample was then centrifuged at 2000x*g* for 5 minutes at 4°C, and pellet was resuspended in cold buffer 2 (DPBS, 1 M sorbitol, 1.5 U zymolase [Zymo E100])). The sample was then incubated for 5 minutes at 30°C, harvested at 2000x*g* for 5 minutes at 4°C, and washed and centrifuged in 500 μL cold 1 M sorbitol without resuspending. Finally, spheroplasts were resuspended in 500 μL cold 1 M sorbitol, 50 μL was taken and harvested at 2000xg for 10 minutes at 4°C. Liquid was removed by pipette, and spheroplast was resuspended in tagmentation buffer according to the ActiveMotif ATAC-seq kit (#53150) method. After adding the tagmentation mix to the sample, we proceeded with the manufacturer’s method starting with the Tagmentation Reaction and Purification section, to tagment samples and amplify sequencing libraries. Prepared ATACseq libraries were sequenced on either an Illumina Novaseq X or an Element AVITI24 at the University of Michigan Advanced Genomics Core. Fastq file quality was checked using FastQC and aligned to the *C. auris* B8441.v3 genome using the Burrows-Wheeler Aligner (BWA) MEM ^66–68^. Bedgraph coverage files were created using Deeptools bamcoverage ^69^. Peaks were identified using MACS3 ^70^. Raw and processed data are deposited in GEO under the accession number GSE336619.

### Library preparation and sequencing

For insertional mutagenesis, *C. auris* AR0382 competent cells were transformed through electroporation as detailed above, using 600 ng of PCR amplified CaNAT from pTO128 using primers oTO668 and oTO1093. Transformed cells were allowed to outgrow in liquid YPD at 30°C for two hours, then nourseothricin stock was added to a final concentration of 200 μg/mL for an additional outgrowth up to 24 hours. Cells were then harvested through centrifugation for genomic DNA extraction.

To prepare libraries, we simultaneously prepared the genomic DNA and the custom U-linker, keeping components on ice. 1 μg of genomic DNA was digested with MseI (NEB #R0525) for three hours at 37°C in a benchtop thermocycler. Digested genomic DNA was then purified using a Zymo Research commercial clean and concentrate kit (Zymo #D4004) using the manufacturer specifications, with the added step of allowing the binding buffer to mix with the sample for three minutes at room temperature to inactivate the restriction enzyme. To prepare the U-linker (oTO2605), 10 μM oligo was incubated in an annealing buffer (100 mM NaCl, 10 mM Tris, pH 8, 1 mM EDTA, pH 8) for 90°C for two minutes, then cooled to 30°C at a ramp rate of -0.1°C/s. The reaction was then snap-cooled in ice and then additional reagents were added to simulate NEB 2.1 buffer (10 nM MgCl2, 100 μg/mL albumen). MseI was then added to the reaction, which was incubated at 37°C for two hours on a benchtop thermocycler. The reaction was purified using a Zymo oligo clean and concentrate kit, eluted with 21 μL of water, and 18 μL of U-linker was mixed with 2 μL 10x annealing buffer. This was heated on a thermocycler to 80°C for two minutes, then cooled to 23°C at a ramp rate of -0.1°C/s. The resulting digested U-linker reaction was kept on ice until ligation.

To ligate the digested U-linker and genomic DNA, we used the NEB quick ligation kit (NEB #M2200) to mix 2.5 μL of U-linker, 9 μL of digested genomic DNA, 12.5 μL of 2x quick ligation buffer, and 1 μL of ligase together, and incubated the reaction at room temperature for ten minutes. The reaction was stopped by adding 1 μL of 0.5 M EDTA. Ligated DNA was purified using Omega Bio-Tek Mag-Bind TotalPure SPRI beads (#M1378) at a 0.9x ratio of beads to reaction volume. We then PCR-amplified ligated DNA with a first round of PCR to add Illumina sequences with the following master mix: 12.5 μL milli-Q water, 10 μL 5x Q5 reaction buffer, 2.5 μL of 10 μM oTO2516, 2.5μL of 10 μM oTO2517, 1 μL of 25mM dNTP mix, 0.5 μL Hot-start Q5 Polymerase, and 1 μL USER enzyme. 30 μL of master mix was added to 20 μL of ligated DNA, for the following reaction: initial incubation at 37°C for USER activity, an initial denaturation at 95°C for 1 minute, then 12 cycles of denaturation (95°C, 15 seconds), annealing (63°C, 15 seconds), and extension (72°C, 1 minute); this was followed by a final extension cycle (72°C, 2 minutes). The product of this reaction was purified using Omega Bio-Tek Mag-Bind TotalPure SPRI beads (#M1378) at a 0.9x ratio of beads to reaction volume. Purified DNA was then added to a reaction master mix (10 μL 5x Q5 reaction buffer, 1 μL of 25mM dNTP mix, 8.5 μL milli-Q water, 0.5 μL Hot-start Q5 Polymerase, 5μL of 10 μM labeled i5 sequencing primer, 5μL of 10 μM labeled i7 sequencing primer) and run on a Bio-Rad T100 Thermocycler for the following reaction: initial incubation at 37°C for USER activity, an initial denaturation at 98°C for 1 minute, then 5 cycles of denaturation (98°C, 10 seconds), annealing (65°C, 40 seconds), and extension (72°C, 40 seconds); this was followed by a final extension cycle (72°C, 3 minutes). This final library was purified using Omega Bio-Tek Mag-Bind TotalPure SPRI beads (#M1378) at a 0.9x ratio of beads to reaction volume. Libraries were sequenced on either an Illumina NovaSeqX or Element AVITI24 platform through the University of Michigan Advanced Genomics Core.

### Library analysis

Library sequencing quality control was performed by the University of Michigan Advanced Genomics Core. Read quality was confirmed using FastQC and sequences containing the end of the NAT cassette were identified by initially aligning to pTO128 using the Burrows-Wheeler Aligner (BWA) MEM with minimum seed length (-k) 19 and a band width (-w) of 2. Soft clipped reads corresponding to the junction between the NAT cassette and *C. auris* genomic DNA were then extracted using extractSoftClipped from SE-MEI (Dpryan, https://github.com/dpryan79/SE-MEI). Extracted sequences were then aligned to *C. auris* GCA_002759435.3_Cand_auris_B8441_V3 assembly using BWA MEM to identify NAT insertion site regions ^66,67^. Insert sites were processed using bedtools to condense regions to single-nucleotide resolution at the junction site, and adjacent insertion sites were merged. Insert motif enrichment analysis was performed using the MEME suite of tools on extracted genomic sequences fifteen bases upstream and downstream of NAT insertion sites ^71^. Analysis of NAT insert regions was performed in R and visualized using ggplot2 (4.0.0) and karyoploteR ^72^. For the permutation analysis, we simulated random redistribution of genic NAT inserts throughout the *C. auris* coding region. After normalizing by gene length, any gene where the distribution of permuted inserts included zero was categorized as unresolvable and removed. For genes with observed inserts, a conservative threshold of at least five inserts less than the lowest value from the permutations was used to estimate essentiality. Genes with zero observed inserts were also categorized as essential. Some genes were masked based on their proximity to highly repetitive telomeric regions and include B9J08_03013-03019 from the repetitive telomeric region of CM076439.1, and B9J08_04428 on the telomeric region of CM076441.1. 5586 remaining total annotated genes were considered for this analysis. Gene Ontology enrichment was performed through the Candida Genome Database ^73^. Raw and processed data are deposited in GEO under the accession number GSE336613.

### Gene Expression analysis

Previously published RNAseq data from Santanta et al. was used in this study ^8^. Raw reads were assessed for quality using FastQC and trimmed to remove poor quality reads or adapter content using Trimmomatic ^68,74^. Trimmed reads were mapped to the *C. auris* B8441 reference transcriptome (NCBI GCA_002759435.3) and quantified using salmon v1.9 ^75^. Read counts normalized to transcripts per million (TPM) were used for the subsequent analysis.

### Keratinocyte Adhesion Assay and Mapping

Keratinocyte adhesion assays were performed as previously described^9^, measuring the degree of adherence of each *C. auris* isolate to human keratinocytes. Mapping was performed on this phenotype with the microbial GWAS tool Hogwash^31^, using default settings and parameters. Mapping was performed on a clade-by-clade basis.

## SUPPLEMENTARY FIGURES AND FIGURE LEGENDS

**Supplementary Figure 1.**
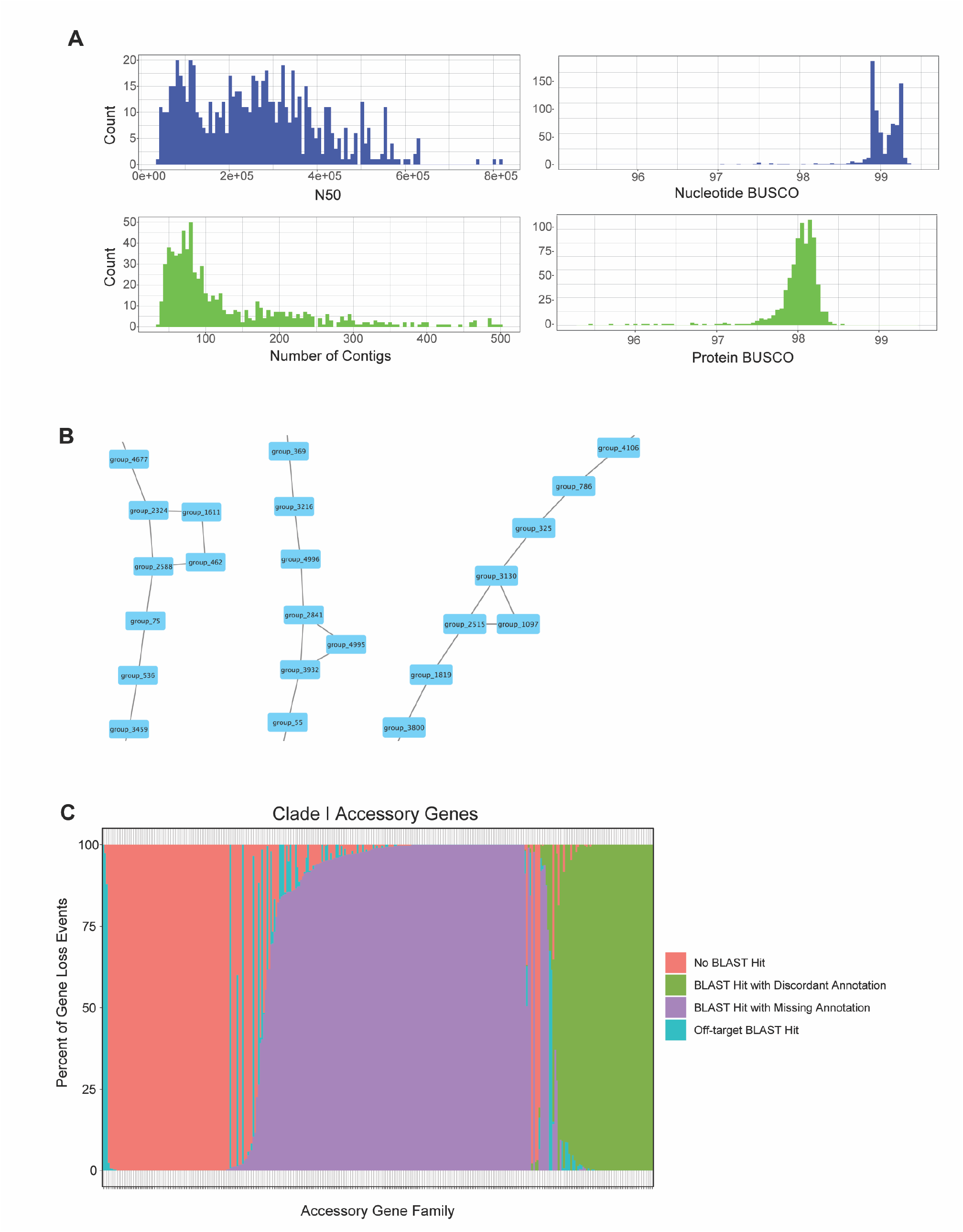
**A,** Metrics of the *de novo* assemblies used as input for pangenome analysis, following all QC steps. Top left: Nucleotide BUSCO score. Bottom left: Protein BUSCO score. Top right: N50. Bottom right: Number of contigs. **B,** Representative example of the pangenome graph produced by Panaroo. Each node represents a single gene family, regardless of how many isolates it appears in. Edges connect gene families if they are adjacent to each other in at least one assembly. This information was used to determine the gene adjacency information for the BLAST validation pipeline. **C,** Representation of BLAST validation outcomes for Clade I accessory genes. Each vertical bar represents a single accessory gene family originally reported by Panaroo, with only gene loss events shown. The outcome of BLAST validation for each of these gene loss events is indicated by color. “No BLAST Hit” indicates concordance between Panaroo and BLAST. “BLAST Hit with Discordant Annotation” indicates that a high-quality BLAST hit was found that overlaps a different, existing gene family in that isolate. “BLAST Hit with Missing Annotation” indicates that a high-quality hit was found, but no annotation was present. “Off-target BLAST Hit” indicates that a high-quality hit was found, but it was not present on the expected scaffold in the expected region.

**Supplementary Figure 2.**
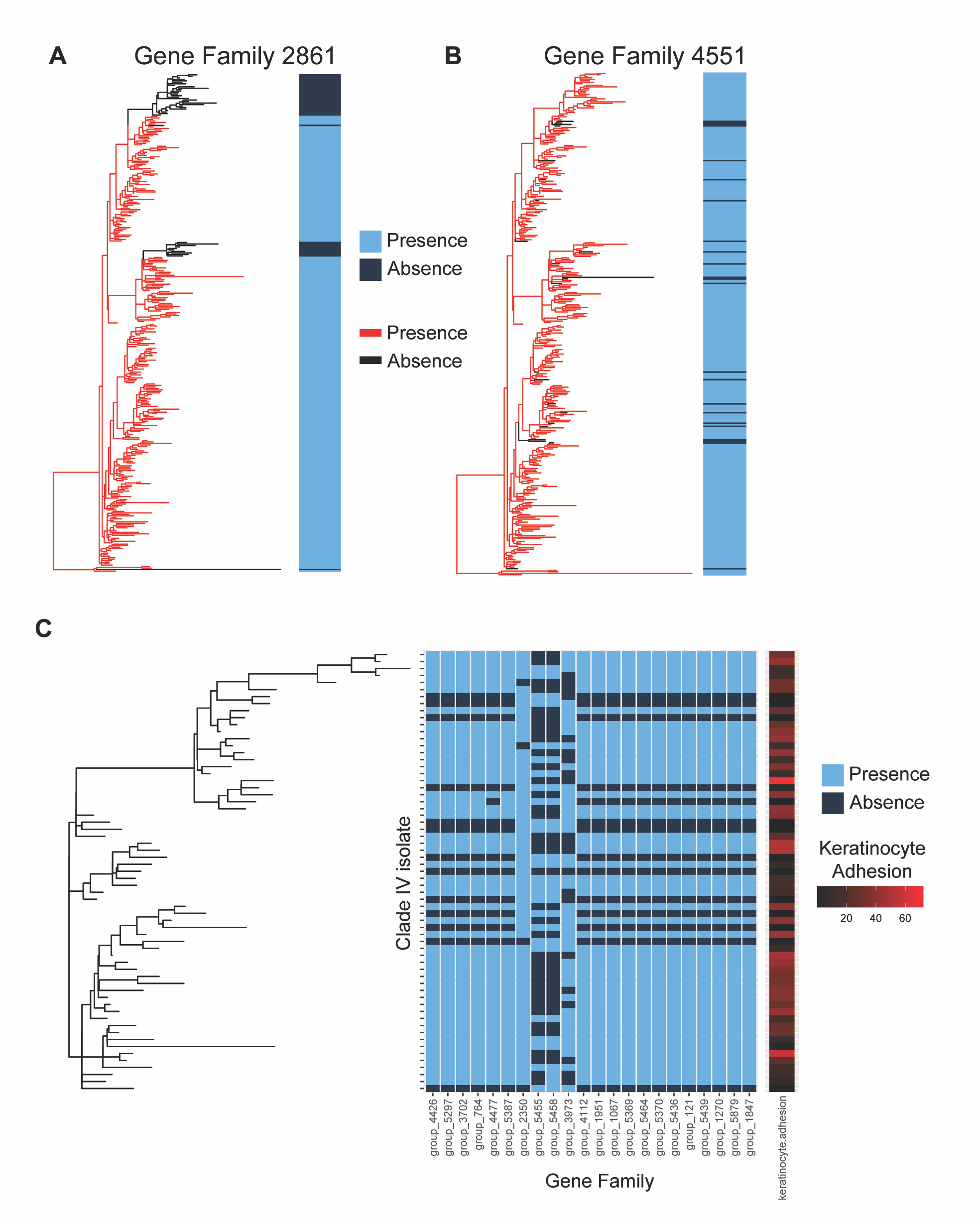
Ancestral state reconstruction. **A,** Representative example of a validated accessory gene from Clade I that shows a strong phylogenetic signal. Most gene loss events cluster tightly on the tree. A total of 40 isolate genomes lack this gene family. **B,** Representative example of a validated accessory gene from Clade I that shows a near-random distribution of gene loss events when compared to the phylogenetic tree. A total of 23 isolate genomes lack this gene family. **C,** Significant mGWAS hits for keratinocyte adhesion in Clade IV isolates. Keratinocytes were used as a model for adhesion to human skin. Each row is a different isolate, and each column is a different gene family with a significant association with keratinocyte adhesion. Gene families are ordered from left to right based on estimated genomic position.

**Supplementary Figure 3.**
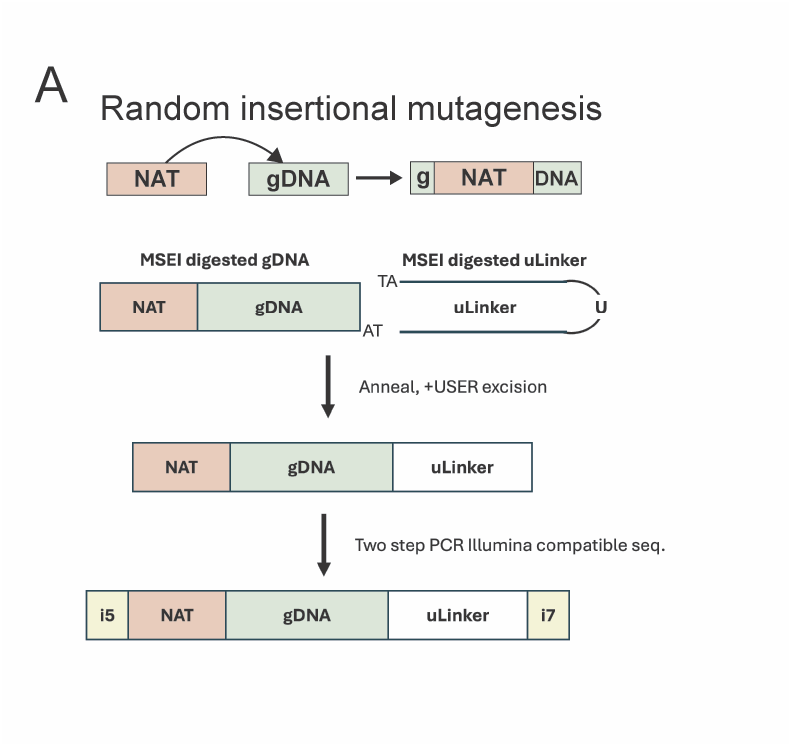
Design for unguided integration mutagenesis using a NAT selection marker. The NAT resistance cassette was transformed into competent *C. auris* cells via electroporation. Genomic DNA was harvested, digested with a MseI restriction enzyme, and ligated to a U-linker to make linear hybrid junction fragments. These fragments were then amplified through two steps of PCR to produce Illumina-compatible sequencing libraries.

**Supplementary Figure 4.**
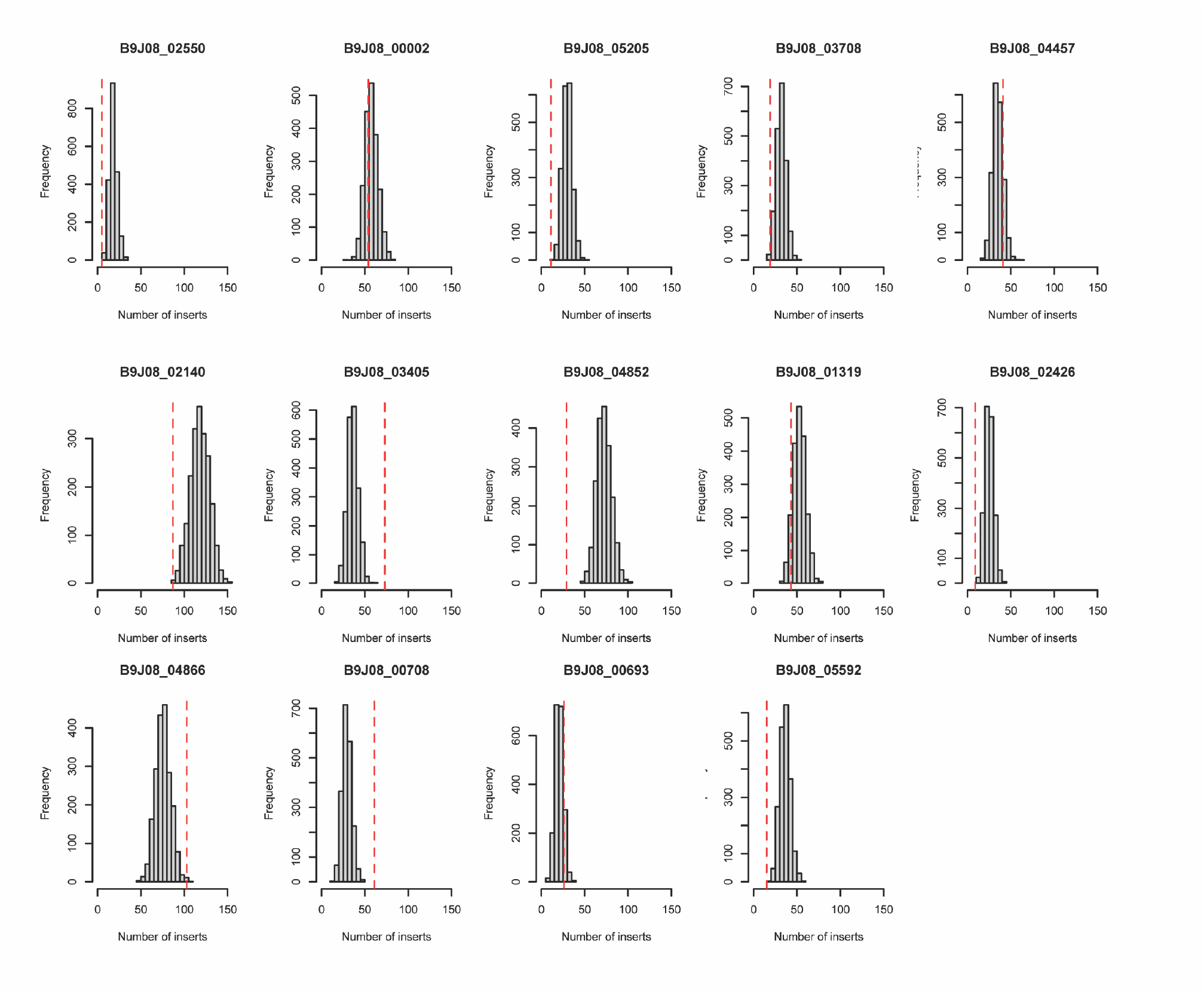
Random permutation simulation analysis distributed the proportion of inserts landing in coding ORFs (∼115k sites) across the gene sample space. Comparing the lower bound of the distribution to our observation (red line) allowed us to further identify ORFs of essential genes.

**Supplementary Figure 5.**
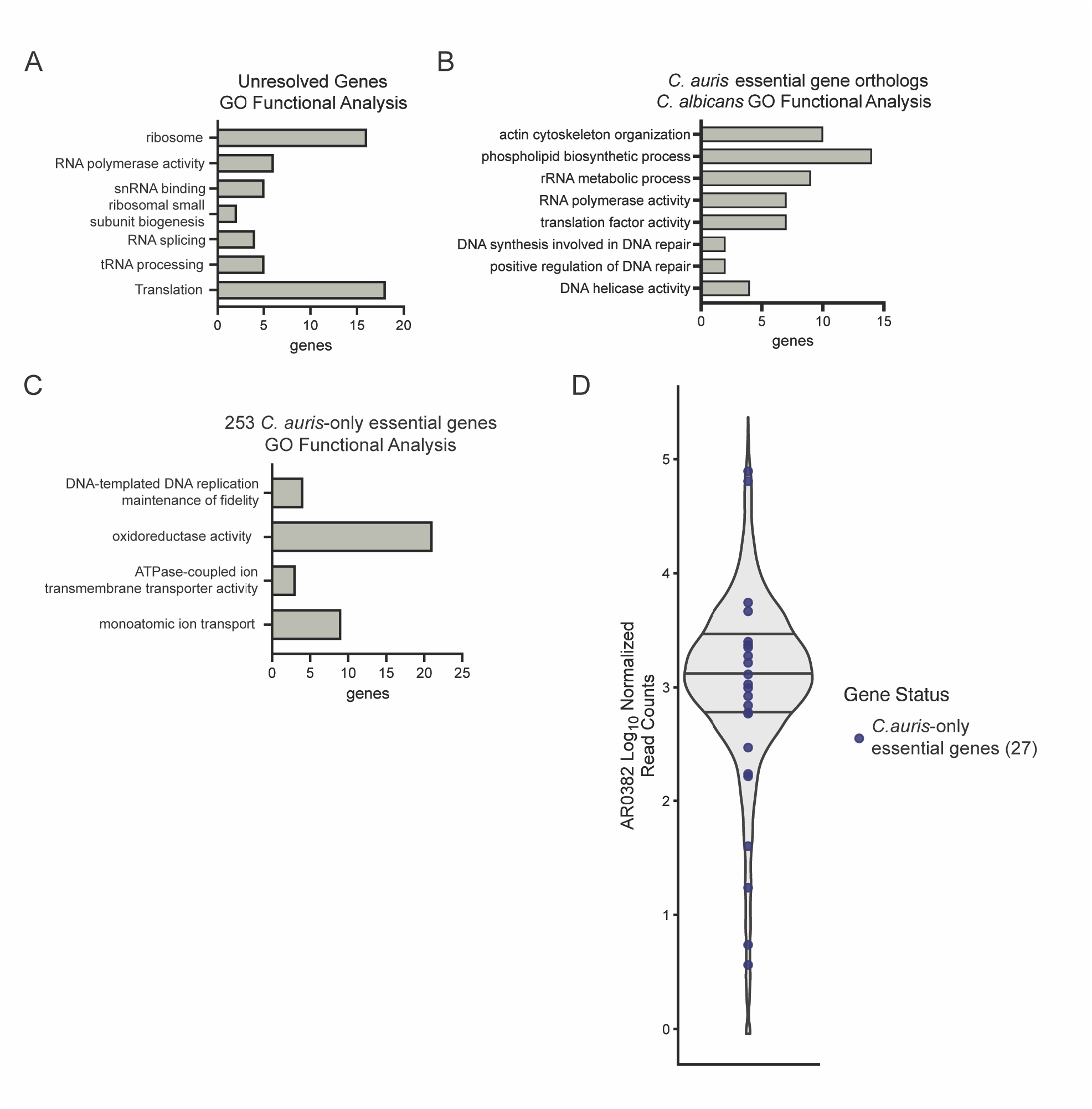
**(A,B,C),** Significantly enriched terms from GO functional analysis of **A,** 219 *C. auris* genes in our defined unresolved category, **B,** 511 *C. auris* essential genes with orthologs in *C. albicans* and **C,** 253 *C.auris* essential genes without essential orthologs in *C. albicans* and *S. cerevisiae*. GO enrichment analysis was performed using the Candida Genome Database GO Term Finder. **D,** RNAseq expression for total *C. auris* genes. Set of 27 *C. auris* essential genes with no orthologs in *C. albicans* and *S. cerevisiae* are labeled in blue. Data from Santana et al. 2023.

**Supplementary Figure 6.**
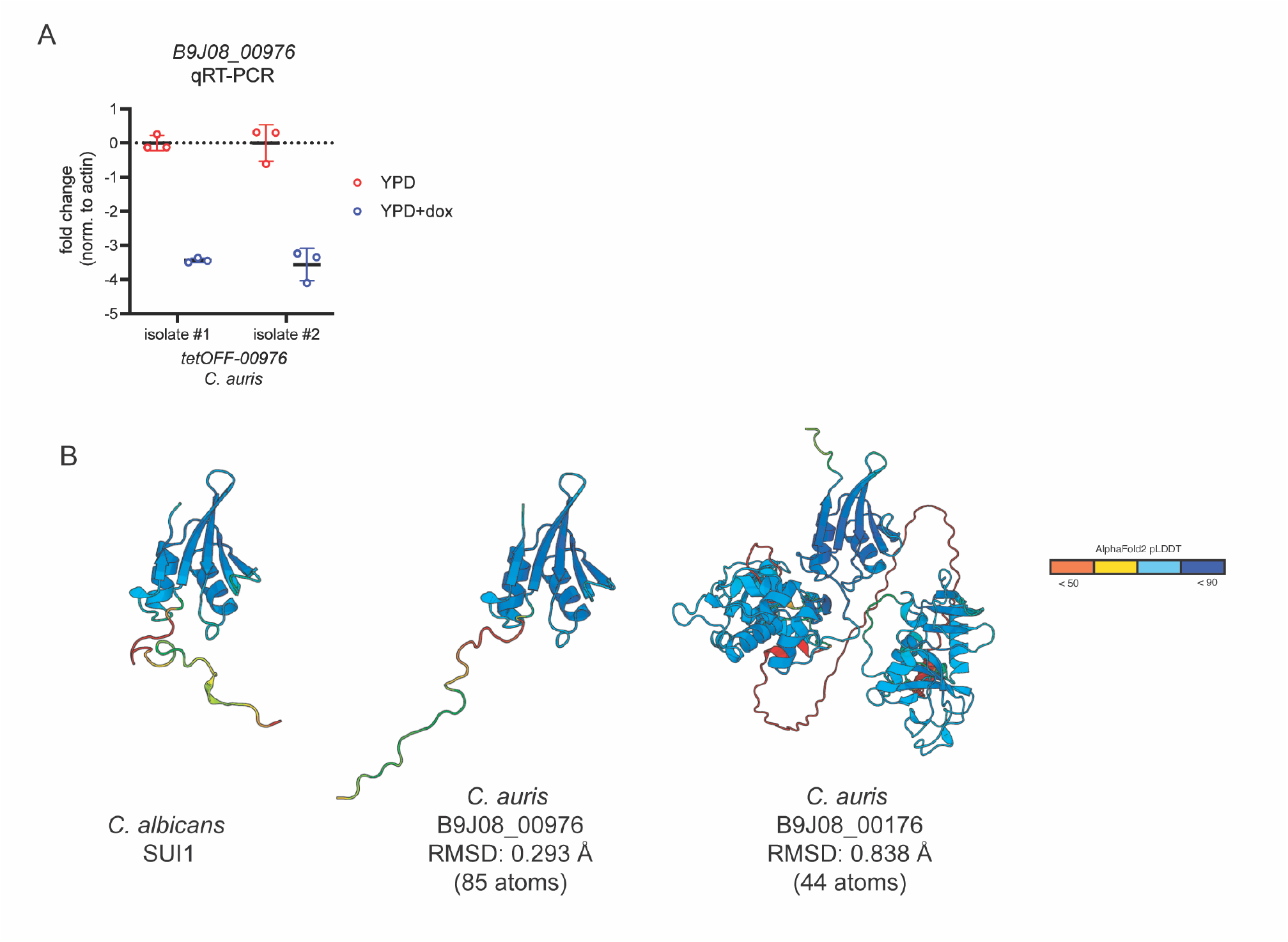
**A,** qRT-PCR for *B9J08_00976* in indicated *C. auris* isolates, with and without 40 μg/mL doxycycline treatment. Data from two independent isolates. **B,** Alphafold predicted structures for the *C. auris* and *C. albicans* Sui1 protein in comparison with B9J08_00176, which has a Sui1-like fold. RMSD over the shared domain is 0.838 Å.

## Supplementary tables

Table S1 Accession numbers and sequencing data used in this study

Table S2 Strains used in this study

Table S3 Oligonucleotides used in this study

Table S4 Plasmids used in this study

Table S5 *ALS4112* deletions among Clade IV isolates from Rush University Medical Center

## Supplementary data

Data S1 Validated pangenome

Data S2 Pangenome CDS names

Data S3 Gene essentiality

