## Supplementary figures and images for "Pangenome and genome-scale essentiality in the emerging fungal pathogen *Candidozyma auris*"

### sup fig 1

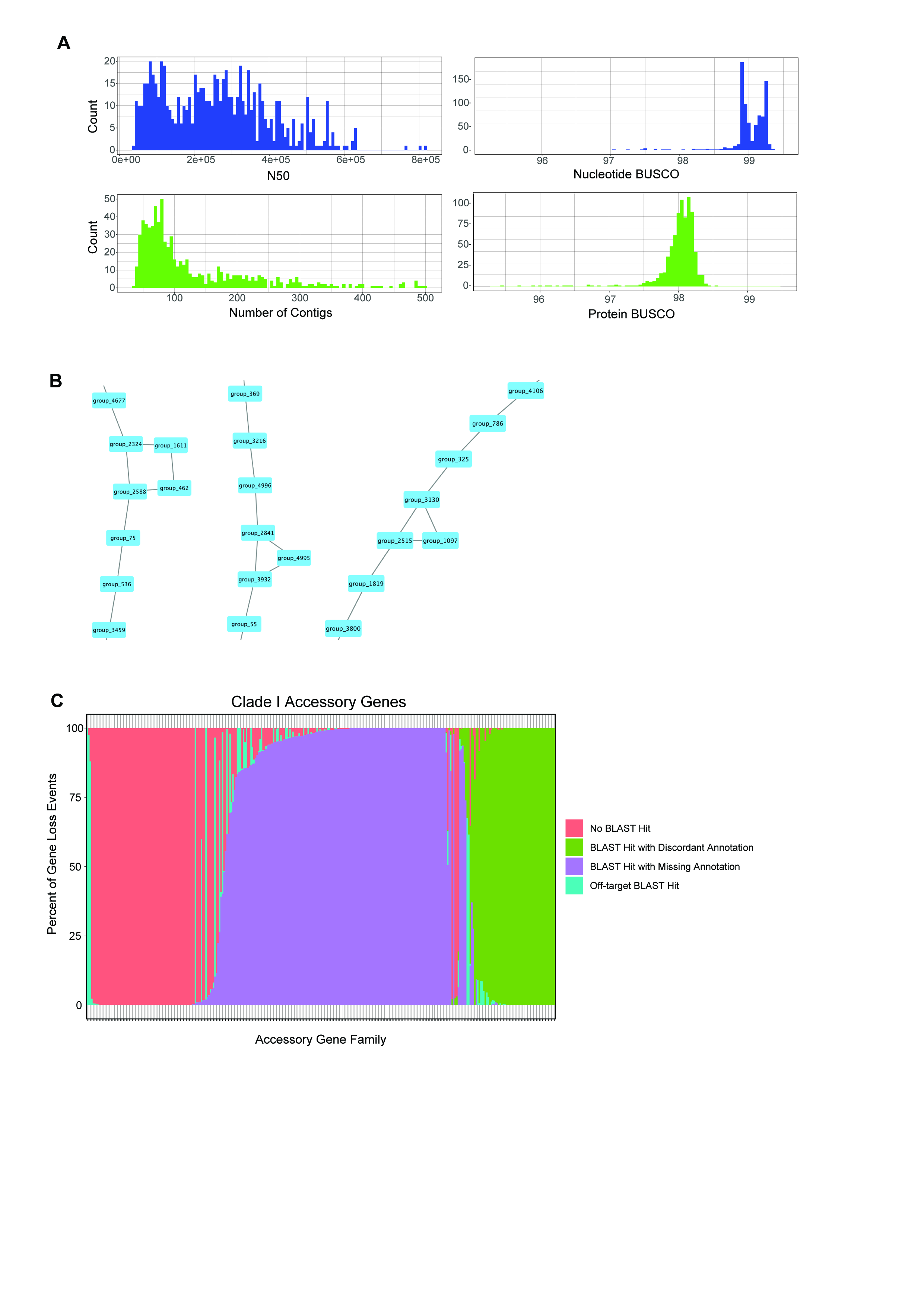

### sup fig 2

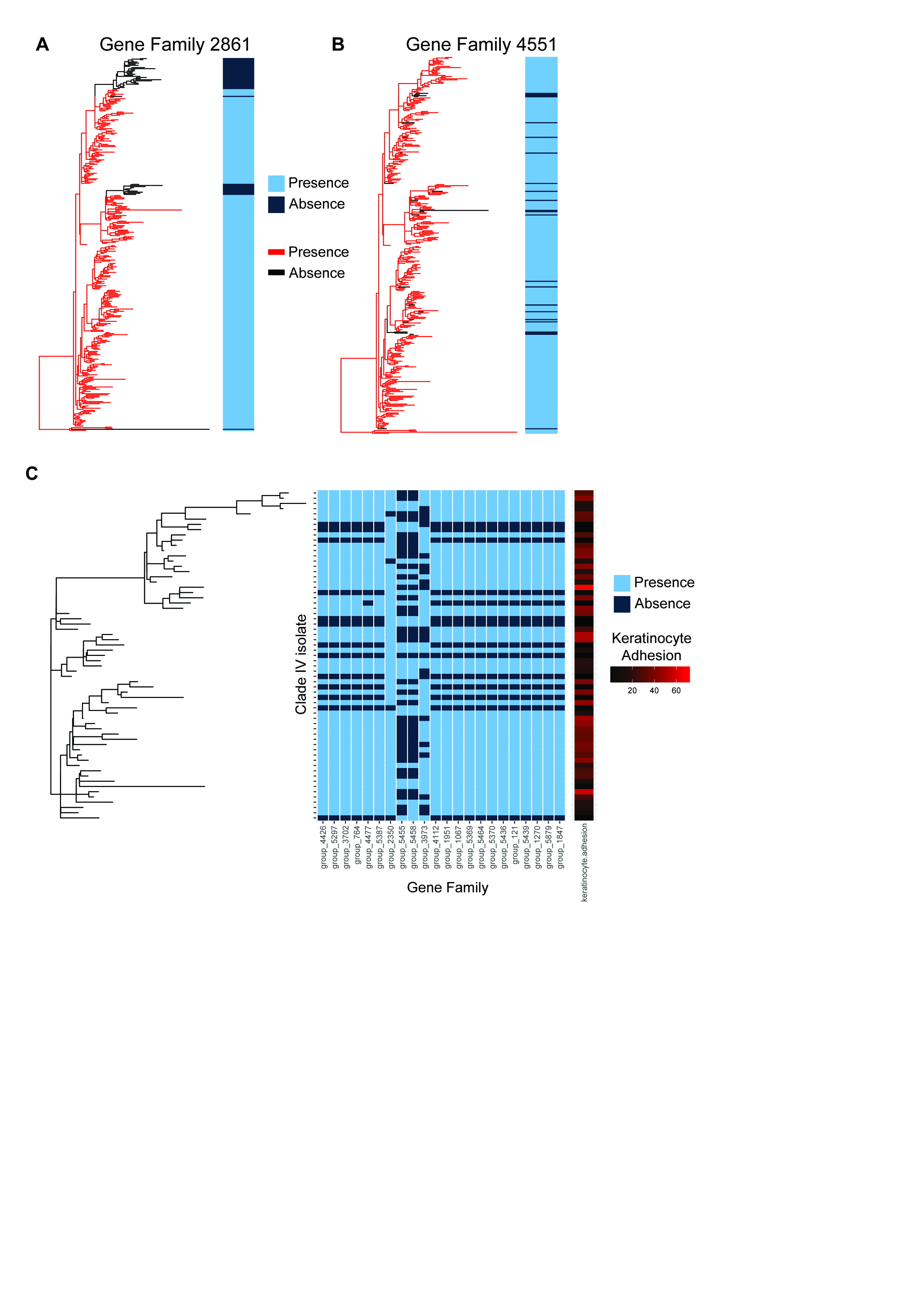

### sup fig 3

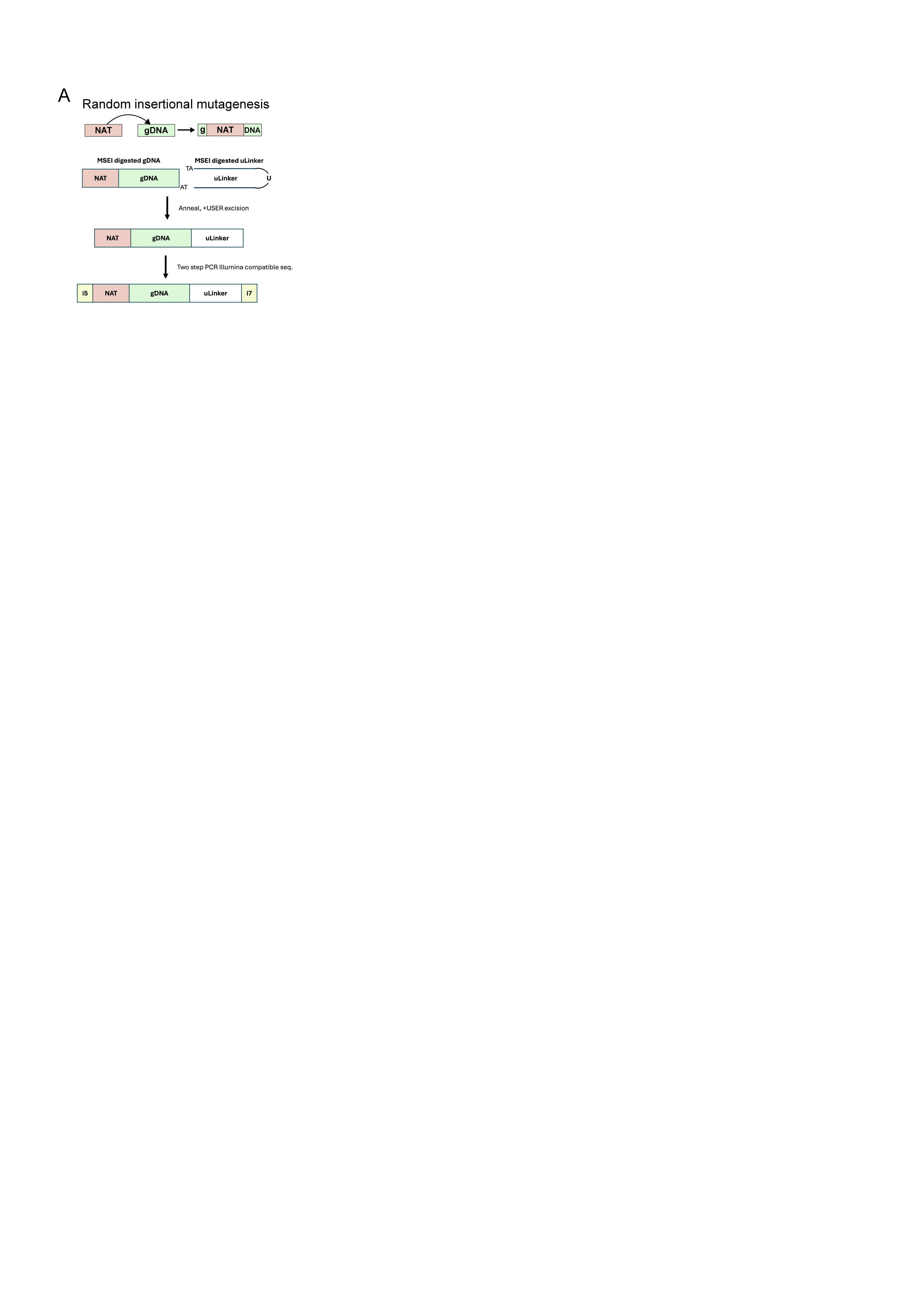

### sup fig 4

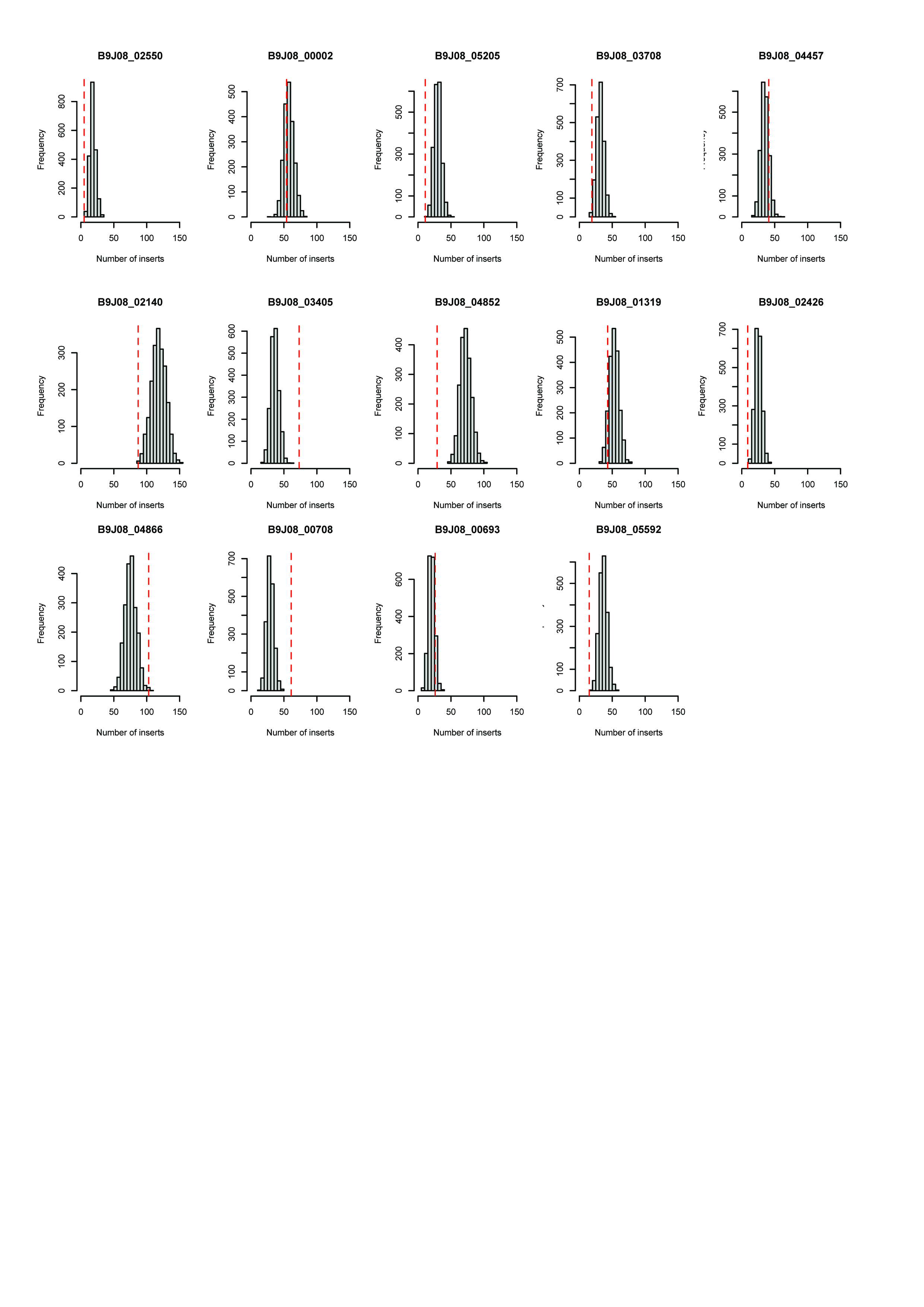

### sup fig 5

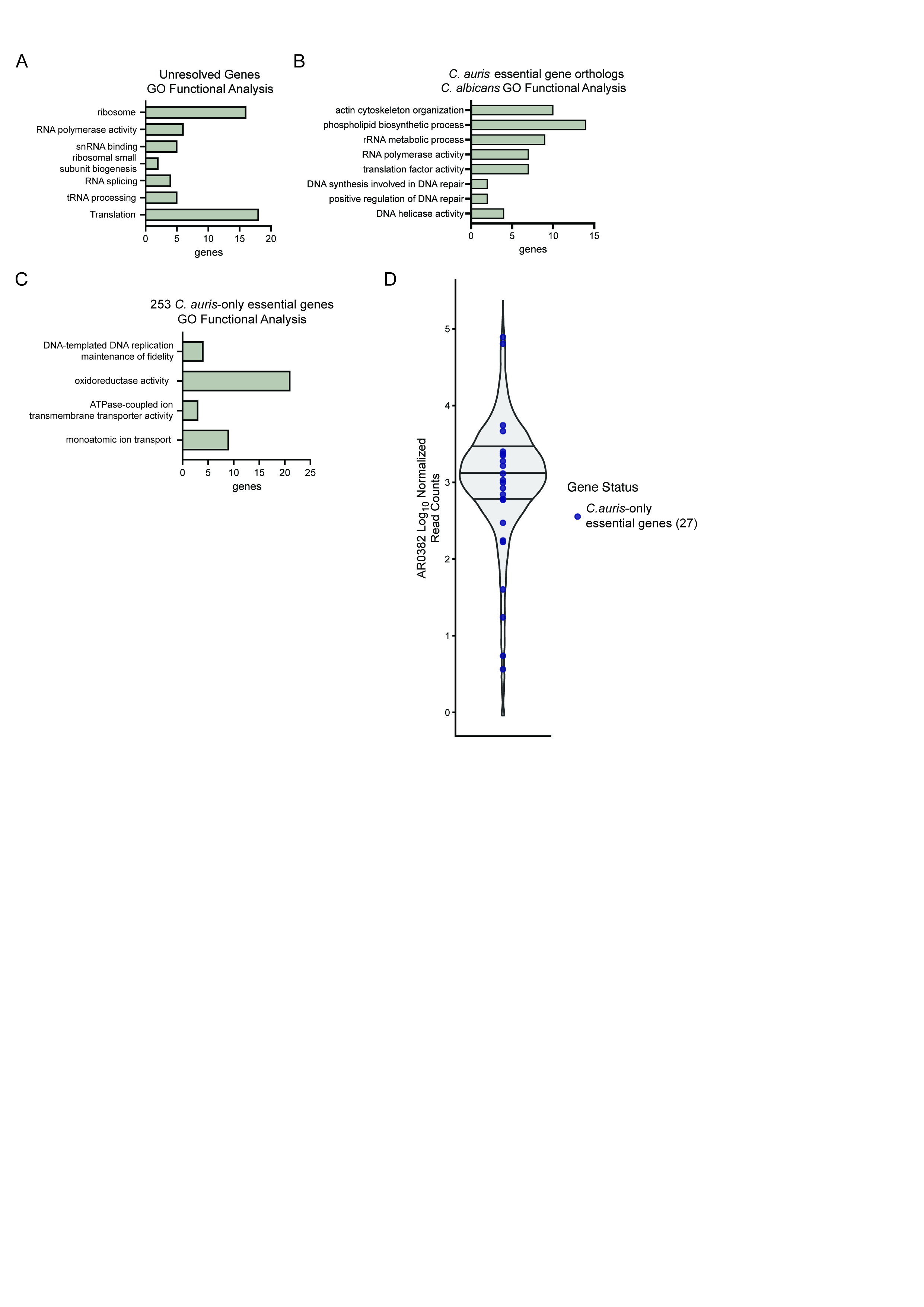

### sup fig 6

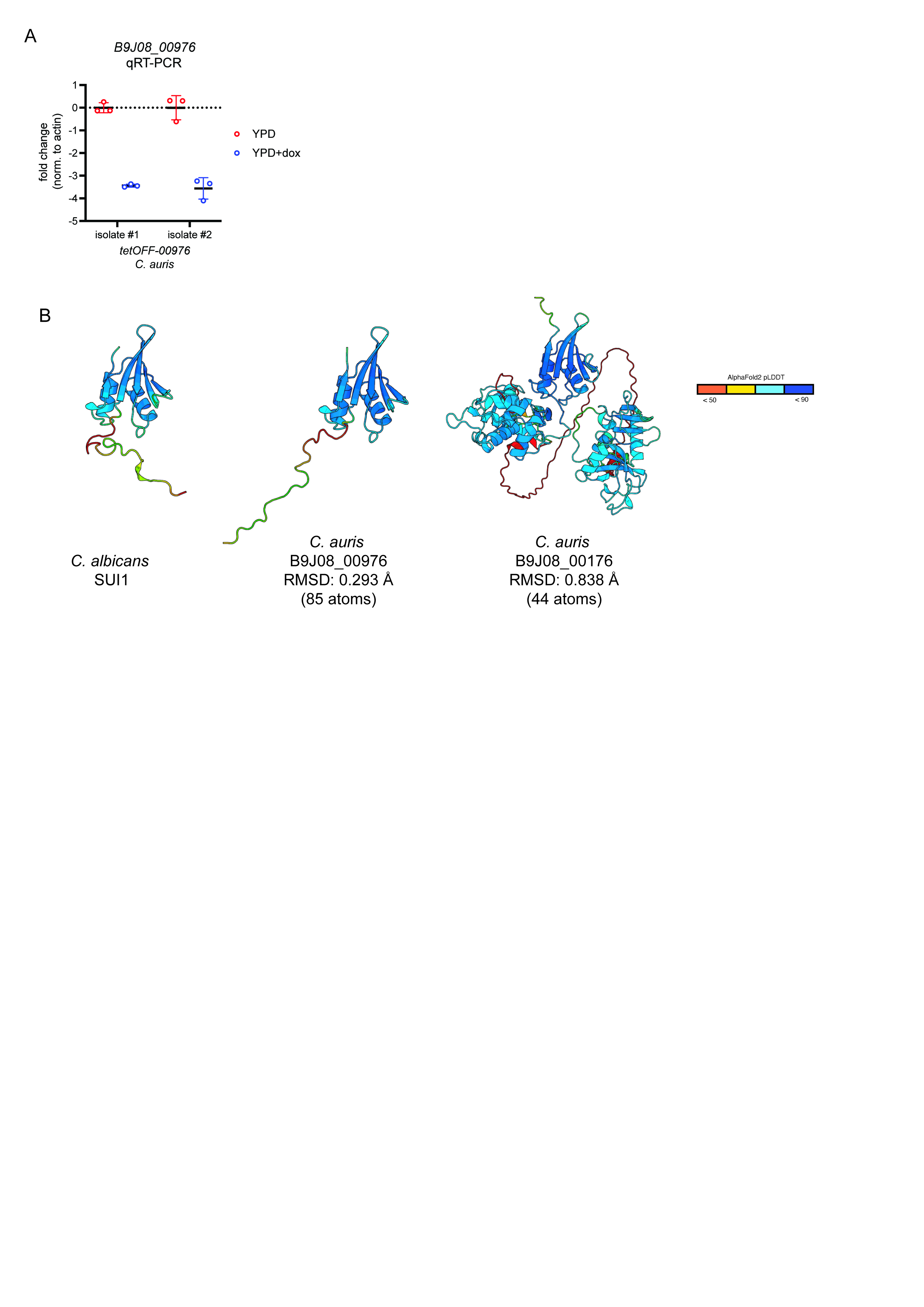
